# Genome-wide characterization of the *MLO* gene family in *Cannabis sativa* reveals two genes as strong candidates for powdery mildew susceptibility

**DOI:** 10.1101/2021.07.16.452661

**Authors:** Noémi Pépin, Francois Olivier Hebert, David L. Joly

**Author notes:** Correspondence: David L. Joly. These authors have contributed equally to this work (Noémi Pépin, Francois Olivier Hebert).

## Abstract

*Cannabis sativa* is increasingly being grown around the world for medicinal, industrial, and recreational purposes. As in all cultivated plants, cannabis is exposed to a wide range of pathogens, including powdery mildew (PM). This fungal disease stresses cannabis plants and reduces flower bud quality, resulting in significant economic losses for licensed producers. The *Mildew Locus O* (*MLO*) gene family encodes plant-specific proteins distributed among conserved clades, of which clades IV and V are known to be involved in susceptibility to PM in monocots and dicots, respectively. In several studies, the inactivation of those genes resulted in durable resistance to the disease. In this study, we identified and characterized the *MLO* gene family members in five different cannabis genomes. Fifteen *Cannabis sativa MLO* (*CsMLO*) genes were manually curated in cannabis, with numbers varying between 14, 17, 19, 18, and 18 for CBDRx, Jamaican Lion female, Jamaican Lion male, Purple Kush, and Finola, respectively (when considering paralogs and incomplete genes). Further analysis of the *CsMLO* genes and their deduced protein sequences revealed that many characteristics of the gene family, such as the presence of 7 transmembrane domains, the MLO functional domain, and particular amino acid positions, were present and well conserved. Phylogenetic analysis of the MLO protein sequences from all five cannabis genomes and other plant species indicated seven distinct clades (I through VII), as reported in other crops. Expression analysis revealed that the *CsMLOs* from clade V, *CsMLO1* and *CsMLO4*, were significantly upregulated following *Golovinomyces ambrosiae* infection, providing preliminary evidence that they could be involved in PM susceptibility. Finally, the examination of variation within *CsMLO1* and *CsMLO4* in 32 cannabis cultivars revealed several amino acid changes, which could affect their function. Altogether, cannabis *MLO* genes were identified and characterized, among which candidates potentially involved in PM susceptibility were noted. The results of this study will lay the foundation for further investigations, such as the functional characterization of clade V MLOs as well as the potential impact of the amino acid changes reported. Those will be useful for breeding purposes in order to develop resistant cultivars.

## 1 Introduction

*Cannabis sativa* is a dicotyledonous plant belonging to the Cannabaceae family, and it is considered a socially and economically important crop as it is increasingly being grown and cultivated around the world. In Canada alone, the sales from cannabis stores in 2020 reached over 2.6 billion dollars (Statistics Canada, 2021), and the number of licensed cultivators, processors, and sellers quadrupled from 2018 to 2020 (Health Canada, 2021). It is used as a source of industrial fiber, seed oil, food, as well as for medicinal, spiritual, and recreational purposes (Small, 2015). As in all cultivated plants, cannabis is exposed to numerous pathogens, and the resulting diseases play a limiting role in its production.

Powdery mildew (PM) is a widespread plant disease caused by ascomycete fungi of the order Erysiphales, for which more than 800 species have been described (Braun and Cook, 2012). They are obligate biotrophs that form invasive structures in epidermal cells for nutrient uptake, called haustoria (Glawe, 2008). These pathogens can infect nearly 10,000 monocotyledonous and dicotyledonous plant species and cause significant damage to crops and ornamental plants (Braun et al., 2002). The PM disease in cannabis, caused by *Golovinomyces ambrosiae* emend. (including *Golovinomyces spadiceus*), has been reported on indoor- and greenhouse-grown plants in Canada and in the United States, where the enclosed conditions provide an ideal environment for the germination and propagation of the fungal spores (Pépin et al., 2018; Szarka et al., 2019; Weldon et al., 2020; Farinas and Peduto Hand, 2020, Wiseman et al., 2021). An analysis of cannabis buds revealed *Golovinomyces* sp. in 79% of tested samples (Thompson et al., 2017), highlighting its ubiquity among licensed producers. The symptoms initially appear as white patches on leaves, and eventually, the mycelia progress to cover the entire leaf surface, the flower bracts, and buds, resulting in stressed and weakened plants, reduced yield, and reduced flower buds quality. Fungicides are widely used to prevent and control this disease in agricultural settings. However, a scarce amount of such products are currently approved by Health Canada as the presence of fungicide residues in the inflorescences raises concerns (Punja, 2021). Besides, they are costly, and fungicide resistance in PM has been observed and documented in other plant species in recent years (Vielba-Fernández et al., 2020). Alternative approaches to managing this disease have been described, such as the use of biological control (e.g., *Bacillus subtilis* strain QST 713), reduced risk products (e.g., potassium bicarbonate, knotweed extract), and physical methods (e.g., de-leafing, irradiation) (Punja, 2021). Nonetheless, some of these methods increase production costs, are labor-intensive, and necessitate further research. Therefore, identifying sources of genetic resistance to PM in cannabis and ultimately breeding or developing resistant cultivars offers the most effective and sustainable approach to controlling PM.

A common strategy used in resistance breeding relies on the exploitation of resistance genes in plants, which encodes for cytoplasmic receptors such as nucleotide-binding leucine-rich repeat proteins or surface receptors such as receptor-like kinases and receptor-like proteins. These immune receptors can detect specific proteins or molecules produced by the pathogen and subsequently induce plant defense responses (Dangl and Jones, 2001). While useful, most resistance genes confer race-specific resistance and are therefore frequently overcome by the emergence of a pathogen’s new virulent race within a few years. An alternative approach in resistance breeding is to exploit susceptibility genes (*S-genes*) in plants. *S-genes* are defined as genes that facilitate infection and support compatibility for a pathogen (van Schie and Takken, 2014). The alteration of such genes can limit the pathogen’s ability to infect the plant and therefore provide a durable type of resistance (van Schie and Takken, 2014). Such PM resistance was initially observed in an X-ray irradiated barley (*Hordeum vulgare*) population in the 1940s (Freisleben and Lein, 1942). It was discovered later that the immunity was attributable to a mutated *S-gene* named *Mildew Locus O* (*MLO*), which was recessively inherited. Complete resistance to all known isolates of PM, caused by *Blumeria graminis* f. sp. *hordei* was conferred in barley by these loss-of-function mutations when present in the homozygous state. This type of resistance has been durable under field conditions and has been used for over forty years in barley breeding programs, without any break in the resistance (Jørgensen, 1992; Büschges et al., 1997; Lyngkjær and Carver, 2000; Piffanelli et al., 2002).

Since discovering *MLO* genes in barley, many *MLO* homologs have been identified in several plant species, especially in monocots and eudicots, as the PM disease affects angiosperms solely. For instance, *MLO* genes were identified in Rosaceae (roses (Kaufmann et al., 2012), apple, peach, strawberry and apricot (Pessina et al., 2014)), Cucurbitaceae (cucumber (Zhou et al., 2013), melon, watermelon, zucchini (Iovieno et al., 2015) and pumpkin (Win et al., 2018)), Solanaceae (tomato (Bai et al., 2008), pepper (Zheng et al., 2013), tobacco, potato and eggplant (Appiano et al., 2015)), Fabaceae (pea (Humphry et al., 2011; Pavan et al., 2011), soybean (Shen et al., 2012; Deshmukh et al., 2014), barrel medic, chickpea, narrow-leaf lupin, peanut, pigeon pea, common bean, mung bean (Rispail and Rubiales, 2016) and lentil (Polanco et al., 2018)), Brassicaceae (thale cress (Devoto et al., 1999; Devoto et al., 2003)), Vitaceae (grapevine (Feechan et al., 2008)), and Poaceae (rice (Liu and Zhu, 2008), wheat (Konishi et al., 2010), sorghum (Singh et al., 2012), maize (Devoto et al., 2003; Kush et al., 2016), foxtail millet (Kush et al., 2016) and stiff brome (Ablazov and Tombuloglu, 2016)). Furthermore, thorough phylogenetic analyses of land plants revealed that *MLO* genes were not only present in monocots and eudicots but also in basal angiosperms, gymnosperms, lycophytes, and bryophytes (Jiwan et al., 2013; Kusch et al., 2016; Shi et al., 2020). Many MLO-like proteins were also identified in algae and other unicellular eukaryotes, suggesting that *MLO* is an ancient eukaryotic protein (Jiwan et al., 2013; Kusch et al., 2016; Shi et al., 2020).

The *MLO* gene family is described as a medium-sized plant-specific gene family, with a varying number of members between 7 in wheat to 39 in soybean, depending on the species (Acedevo-Garcia et al., 2014). The resulting MLO proteins are characterized by the presence of seven transmembrane domains integral to the plasma membrane with an extracellular N-terminus and an intracellular C-terminus (Devoto et al., 1999). They are also characterized by the presence of a calmodulin-binding domain in the C-terminal region that is likely implicated in sensing calcium influx and mediating various signaling cascades (Kim et al., 2002a; Kim et al., 2002b). MLO protein sequences identified across land plants also possess several highly conserved amino acids, some of which have been deemed essential for the structure, functionality, and stability of the protein (Devoto et al., 2003; Elliott et al., 2005; Reinstädler et al., 2010; Kusch et al., 2016). Mutations in these residues could affect the accumulation, maturation, and function of the protein and are therefore attractive targets for breeding programs (Elliott et al., 2005; Reinstädler et al., 2010).

Throughout land plant evolution, the MLO protein family diversified into subfamilies, or clades, which have been demonstrated in several phylogenetic analyses. MLO proteins are usually grouped into seven defined clades (I to VII), among which clades IV and V appear to host MLO proteins associated with PM susceptibility in monocots and dicots, respectively (Acevedo-Garcia et al., 2014; Kusch et al., 2016). It has been documented in many species, such as barley, tomato, and apple, that *MLO* genes from these two clades (IV and V in monocots and dicots, respectively) are up-regulated upon PM infection (Piffanelli et al., 2002; Zheng et al., 2013; Pessina et al., 2014). It has also been demonstrated that the overexpression of these genes results in enhanced susceptibility to the pathogen (Zheng et al., 2013). Furthermore, the inactivation of these genes in many species by gene silencing, genome editing, or TILLING has resulted in increased or complete resistance to PM (Wang et al., 2014; Acevedo-Garcia et al., 2017; Nekrasov et al., 2017; Ingvardsen et al., 2019; Wan et al., 2020). Besides the implication of clade V and IV MLOs in PM susceptibility, recent studies have suggested that *MLO* genes from other clades are implicated in various physiological and developmental processes. For example, it was demonstrated that in *Arabidopsis thaliana*, *AtMLO4* and *AtMLO11* from clade I are involved in root thigmomorphogenesis (Chen et al., 2009; Bidzinski et al., 2014), while *AtMLO7* from clade III is involved in pollen tube reception by the embryo sac (Kessler et al., 2010). In rice (*Oryza sativa*), *OsMLO12* from clade III mediates pollen hydration (Yi et al., 2014). Interestingly, barley *HvMLO1* has been shown to differentially regulate the establishment of mutualistic interactions with the endophyte *Serendipita indica* and the arbuscular mycorrhizal fungus *Funneliformis mosseae* (Hilbert et al., 2020). Indeed, another study clearly showed barley *HvMLO1*, wheat *TaMLO1,* and barrel medic *MtMLO8* from clade IV to be involved in the establishment of symbiotic relationships with beneficial mycorrhizal fungi (Jacott et al., 2020). These findings suggest that powdery mildews might have appropriated and exploited these genes as an entryway to successful pathogenic colonization (Jacott et al., 2020). However, in pea, no evidence was found for the implication of *PsMLO1*, a clade V gene, in the establishment of relationships with mycorrhizal and rhizobial symbionts (Humphry et al., 2011).

Although *MLO* genes have been studied in many monocot and dicot species, they have only been preliminarily studied in cannabis (McKernan et al., 2020). The growing interest in cannabis research has led to the publication of several genomes in recent years, thus providing an opportunity to conduct a comprehensive analysis of the *MLO* gene family in cannabis. In this study, we manually curated and characterized the members of the *MLO* gene family in cannabis from five different available genomes: Purple Kush and Finola (Laverty et al., 2019), CBDRx (Grassa et al., 2021), and Jamaican Lion (female, McKernan et al., 2018; male, McKernan et al., 2020). Through phylogenetic analysis, we identified candidate *MLO* genes likely to be involved in PM susceptibility in cannabis, observed their subcellular localization by confocal microscopy, and monitored their expression profile in cannabis leaves during infection. We also searched for potential naturally occurring resistant mutants by investigating amino acid changes in 32 cultivars. A better understanding of cannabis *MLOs* offers enormous opportunities to breed PM-resistant cultivars and develop new control methods, thereby increasing productivity and yield.

## 2 Materials and methods

### 2.1 *In silico* identification and manual curation of the cannabis *MLO* genes

Cannabis *MLO* genes in CBDRx were initially identified (also named ‘cs10’, NCBI accession GCA_900626175.2) using TBLASTN from the BLAST+ suite (Camacho et al., 2009) with *Arabidopsis thaliana* amino acid sequences as queries (*AtMLO1*-*AtMLO15*, NCBI accession numbers: NP_192169.1, NP_172598.1, NP_566879.1, NP_563882.1, NP_180923.1, NP_176350.1, NP_179335.3, NP_565416.1, NP_174980.3, NP_201398.1, NP_200187.1, NP_565902.1, NP_567697.1, NP_564257.1, NP_181939.1). In parallel, all official gene models were extracted from the NCBI CBDRx annotation report (NCBI Annotation Release 100, https://www.ncbi.nlm.nih.gov/genome/annotation_euk/Cannabis_sativa/100/) with an InterPro (IPR004326) and/or Pfam (PF03094) identification number related to the *MLO* gene family (Blum et al., 2021; Mistry et al., 2021). These two sequence datasets were merged together, and multiple sequence alignments were performed using MUSCLE v.3.8.31 (Edgar, 2004) with the genomic and the amino acid versions of each *MLO* gene model. Only unique sequences were kept, and all *MLO* gene models that were retained but incomplete were further manually curated. The full genomic sequences were aligned and manually compared with their corresponding full-length mRNA transcripts using BLASTN from the BLAST+ suite (Camacho et al., 2009). Each gene was characterized based on total length, chromosome localization, strand, START and STOP positions, as well as number and size of exons and introns (**Figure 1**, **Table 1**). The resulting streamlined and manually curated *MLO* gene models for CBDRx were considered the definitive reference set for this genome.

**Figure 1.**
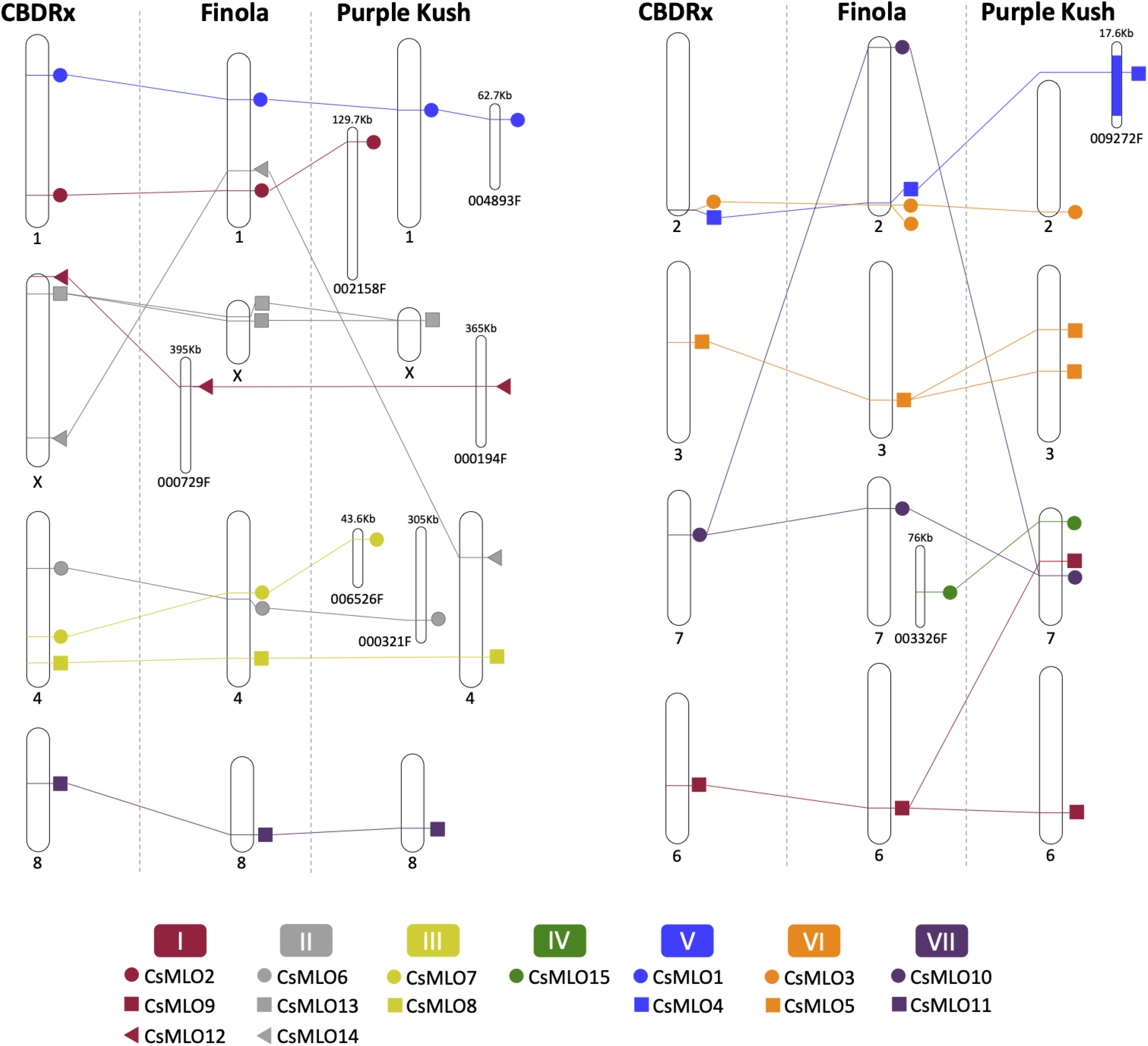
Physical map of *MLO* genes identified in the chromosome-level assemblies of CBDRx, Finola and Purple Kush. Each thick white bar represents a chromosome, with its size exactly proportional to the actual size in base pairs. Numbers below the chromosomes represent the “unified” chromosome numbers according to the CBDRx reference and the orthology analysis that we performed on all genomes. Colors indicate MLO clades from I to VII. In each CsMLO clade, each gene is designated by a different symbol (circle, square or triangle). Homology relationships between the three genomes are displayed with solid lines of colors corresponding to each MLO clade. The *MLO* gene from clade IV (herein named *CsMLO15*) is absent from the CBDRx assembly. Chromosomes 5 and 9 do not carry any *CsMLO* gene and are thus not represented in this figure.

**Table 1.** Members of the *CsMLO* gene family as predicted and manually curated in the CBDRx genome.

| Gene | Clade | Chr. | Strand | Start position | End position | Genomic length (bp) | Exons | Protein length (aa) | TMs<sup>b</sup> | Subcellular localization<sup>c</sup> | Conserved aa/30<sup>d</sup> | Conserved aa/58<sup>e</sup> |
| --- | --- | --- | --- | --- | --- | --- | --- | --- | --- | --- | --- | --- |
| *CsMLO1-CBDRx* | V | 1 | + | 21,631,499 | 21,636,477 | 4,979 | 15 | 555 | 7 | Cell membrane | 28 | 57 |
| *CsMLO2-CBDRx* | I | 1 | - | 84,421,095 | 84,415,528 | 5,568 | 15 | 551 | 7 | Cell membrane | 30 | 58 |
| *CsMLO3-CBDRx* | VI | 2 | - | 92,590,377 | 92,584,053 | 6,325 | 15 | 547 | 7 | Cell membrane | 25 | 52 |
| *CsMLO4-CBDRx* | V | 2 | + | 92,612,089 | 92,622,483 | 10,395 | 15 | 631 | 7 | Cell membrane | 30 | 56 |
| *CsMLO5-CBDRx* | VI | 3 | - | 42,556,462 | 42,507,454 | 49,009 | 15 | 515 | 7 | Cell membrane | 30 | 58 |
| *CsMLO6-CBDRx* | II | 4 | - | 29,809,460 | 29,799,903 | 9,558 | 14 | 520 | 7 | Cell membrane | 30 | 57 |
| *CsMLO7-CBDRx* | III | 4 | - | 65,440,972 | 65,435,678 | 5,295 | 15 | 549 | 7 | Cell membrane | 29 | 57 |
| *CsMLO8-CBDRx* | III | 4 | + | 79,070,668 | 79,074,427 | 3,760 | 15 | 542 | 7 | Cell membrane | 30 | 57 |
| *CsMLO9-CBDRx* | I | 6 | + | 49,150,225 | 49,156,766 | 6,542 | 15 | 568 | 7 | Cell membrane | 30 | 57 |
| *CsMLO10-CBDRx* | VII | 7 | - | 23,870,119 | 23,830,282 | 39,838 | 14 | 535 | 7 | Cell membrane | 30 | 57 |
| *CsMLO11-CBDRx* | VII | 8 | - | 51,420,983 | 51,416,370 | 4,614 | 14 | 583 | 7 | Cell membrane | 30 | 58 |
| *CsMLO12-CBDRx* | I | X | + | 443,359 | 447,357 | 3,999 | 14 | 558 | 7 | Cell membrane | 29 | 55 |
| *CsMLO13-CBDRx* | II | X | + | 10,840,654 | 10,844,220 | 3,567 | 14 | 508 | 7 | Cell membrane | 30 | 58 |
| *CsMLO14-CBDRx* | II | X | + | 89,459,459 | 89,463,372 | 3,914 | 12 | 516 | 7 | Cell membrane | 30 | 56 |
| *CsMLO15-CBDRx*<sup>a</sup> | IV | N/A | N/A | N/A | N/A | N/A | N/A | N/A | N/A | N/A | N/A | N/A |

<sup>a</sup> *CsMLO15-CBDRx* is not predicted in the CBDRx genome, as compared to the other four genomes analyzed.
<sup>b</sup> Number of transmembrane domains (TMs) in the predicted protein, as determined by the CCTOP online prediction server (Dobson et al., 2015).
<sup>c</sup> Subcellular localization as predicted by three online prediction servers: Plant-mPLoc (Chou and Shen, 2010), YLoc-HighRes (Briesmeister et al., 2010), and DeepLoc 1.0 (Armenteros et al., 2017).
<sup>d</sup> Number of conserved amino acids out of the 30 identified in Elliott et al., 2005.
<sup>e</sup> Number of conserved amino acids out of the 58 identified in Kusch et al., 2016.

The reference set of CBDRx *MLO* genes were used as queries to search for the presence of homologs in four other cannabis genomes (Purple Kush - GCA_000230575.5, Finola - GCA_003417725.2, Jamaican Lion female - GCA_012923435.1, Jamaican Lion male - GCA_013030025.1), using TBLASTN. Manual curation of each set of *MLO* genes was performed in each of these genomes, using the same approach described previously for CBDRx. In the process, several frameshifts (mostly small insertions and deletions) were noted and manually corrected in the coding sequence of multiple *MLOs* in Finola (13 genes) and Purple Kush (7 genes). None of the homologs for these genes showed any frameshift in any other genome. All the frameshifts identified in Finola and Purple Kush were thus examined by comparing their respective coding sequence with available transcriptomic data from the CanSat3 assembly project (http://genome.ccbr.utoronto.ca/downloads.html), using BLASTN. Based on evidence from mRNA sequences, all these frameshifts were manually corrected. All of the manually curated *MLO* genes for all five cannabis genomes were ultimately considered as our reference and final set of *Cannabis sativa MLO* (*CsMLO*) genes (**Table 1, Supplementary Tables S1-S5, Supplementary Files S1-S3**). The structure of our final *CsMLO* gene models was compared to their respective genome annotation available on NCBI (CBDRx, Jamaican Lion male and female) (**Supplementary Tables S1, S2 and S3**).

Chromosomal localization of each manually curated *CsMLO* gene in the chromosome-level genome assemblies, i.e. CBDRx, Finola, and Purple Kush, was displayed on a chromosomal map (**Figure 2**) with the R package RIdeogram (Hao et al., 2020). Chromosome numbers in Finola and Purple Kush were standardized to match the official chromosome numbers in the CBDRx reference genome, using whole genome alignments on the D-Genies platform (http://dgenies.toulouse.inra.fr/). The SVG output from RIdeogram was aesthetically modified and adapted using Microsoft © PowerPoint v.16.49 (Microsoft Corporation, Redmond, Washington, U.S.). This analysis served as a preliminary assessment of synteny for *CsMLO* genes.

**Figure 2.**
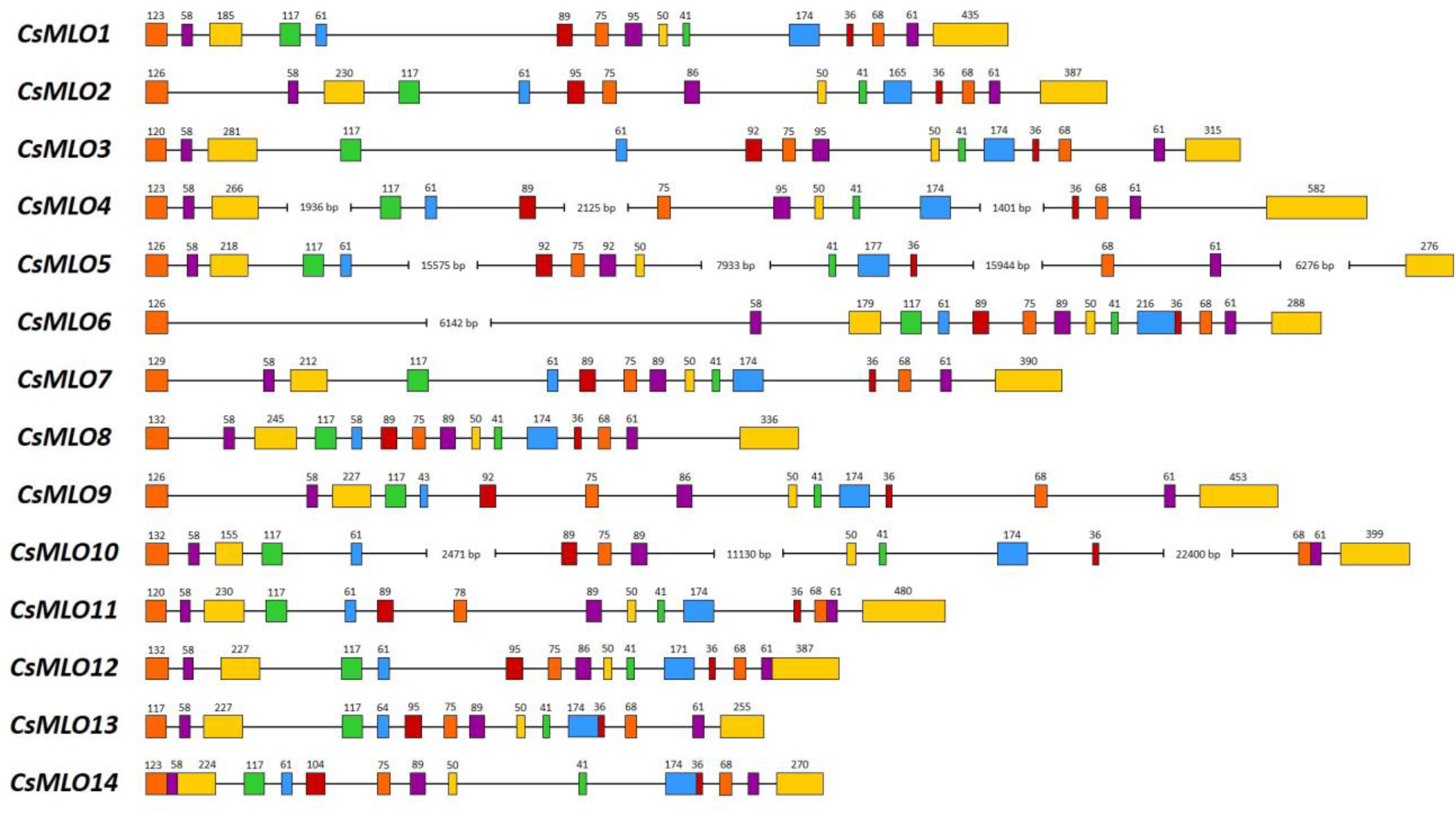
Intron-exon organization of the 14 manually curated *CsMLO* genes identified in the CBDRx genome. Exons are shown as rectangles and introns as lines. The exon color code simply allows demonstrating exon conservation across all sequences. The numbers above exons indicate the exon’s length (bp). Note that the STOP codon (3 bp) is included in the last exon’s length. Sequences exhibiting one or several large introns are severed where necessary (i.e. *CsMLO4*, *CsMLO5*, *CsMLO6* and *CsMLO10*) and are therefore not drawn to scale.

### 2.2 Gene and protein characterization

The protein sequences from the five genomes were analyzed through several online prediction servers in order to identify functional domains (InterProScan, Jones et al., 2014), transmembrane domains (CCTOP, Dobson et al., 2015), subcellular localizations (Plant-mPLoc, Chou and Shen, 2010; DeepLoc 1.0, Almagro Armenteros et al., 2017; YLoc-HighRes, Briesemeister et al., 2010), signal peptide (SignalP 5.0, Almagro Armenteros et al., 2019a), calmodulin-binding domains (CaMELS, Abbasi et al., 2017) as well as mitochondrial, chloroplast, and thylakoid luminal transit peptide (TargetP 2.0, Almagro Armenteros et al., 2019b). Conserved amino acids described in Elliott et al., 2005 (30 invariant amino acids) and Kusch et al., 2016 (58 highly conserved amino acids) were screened in all *CsMLO* sequences using our final protein alignment (**Supplementary File S3**). Our manually curated *MLO* gene models in the five cannabis genomes were also screened for conserved *cis*-acting elements in the promoter regions. A homemade Python script (v.3.7.3, Van Rossum and Drake, 2009) was used to extract a 2 kb upstream region of each *CsMLO* gene, and these promoter regions were used as search queries in the plantCARE database (Lescot et al., 2002).

### 2.3 Phylogenetic analysis

Amino acid sequences for all manually curated CsMLO from CBDRx, Jamaican Lion female and male, Purple Kush and Finola were aligned together with MLO sequences previously identified in *Arabidopsis thaliana* (AtMLO, as indicated in Devoto et al., 2003), *Vitis vinifera* (grapevine: VvMLO, as indicated in Feechan et al., 2008), *Prunus persica* (peach: PpMLO, as indicated in Pessina et al., 2014), *Hordeum vulgare* (barley: HvMLO, as indicated in Kusch et al., 2016) and *Zea mays* (maize: ZmMLO, as indicated in Kusch et al., 2016). *Chlamydomonas reinhardtii* MLO (XP_001689918) was used as an outgroup. Alignment of protein sequences was performed using MAFFT v7.407_1 (Katoh and Standley, 2013) with default parameters within NGPhylogeny.fr (Lemoine et al., 2019) and used to construct phylogenetic trees. A first tree was constructed using PhyML+SMS v1.8.1_1 (Lefort et al., 2017) with default parameters within NGPhylogeny.fr (**Figure 3**), and a second tree was constructed using MrBayes v3.2.6_1 (Huelsenbeck and Ronquist, 2001) with default parameters within NGPhylogeny.fr (**Supplementary Figure S1**). The trees were interpreted and visualized using iTOL (Letunic and Bork, 2019). All MLO proteins identified were classified into clades based on previous phylogenetic analyses (Kusch et al., 2016).

**Figure 3.**
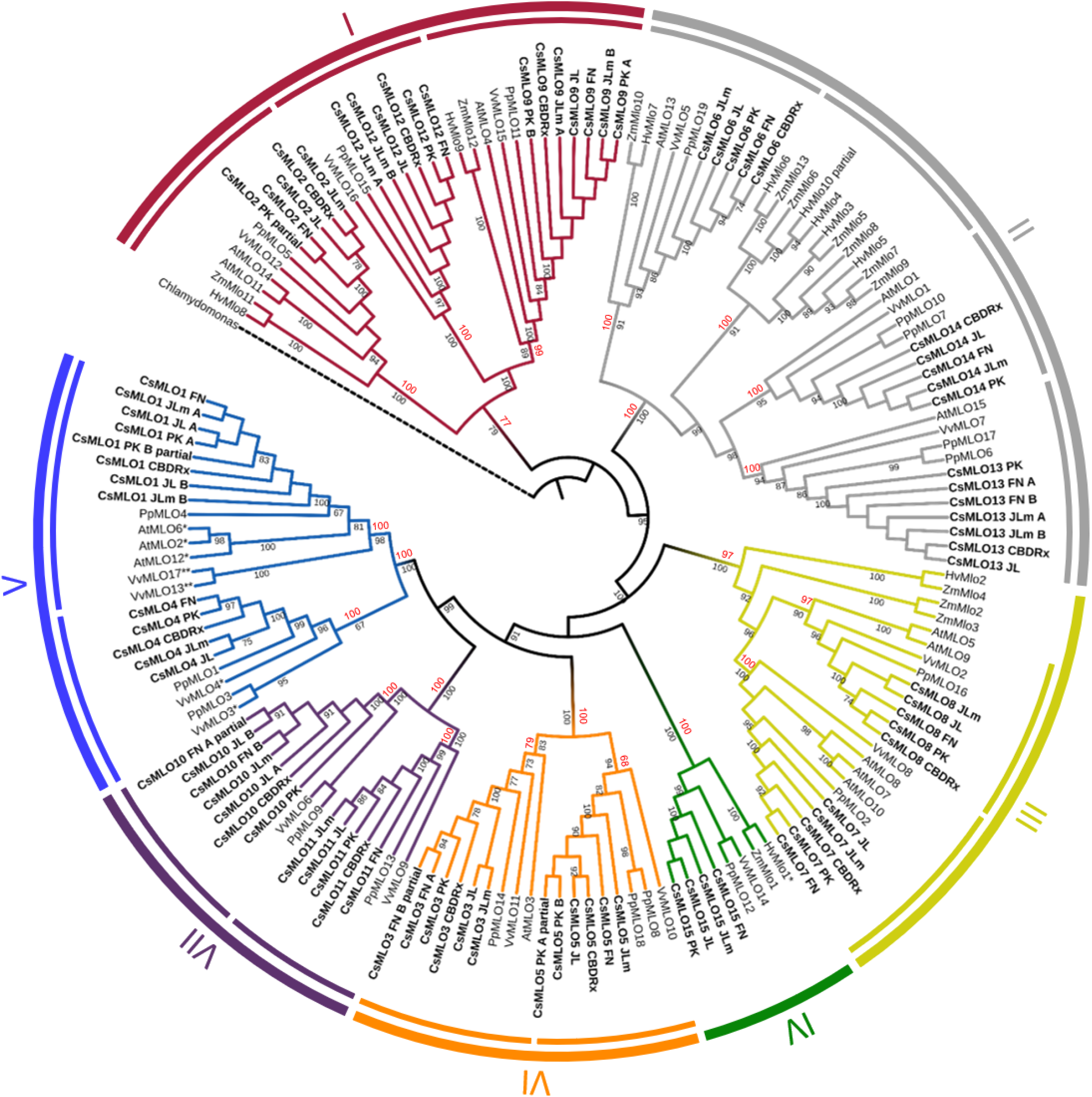
Phylogenetic relationships of CsMLOs based on Maximum Likelihood analysis. Phylogenetic tree of manually curated CsMLO proteins (bold) with MLO proteins from selected species (*Arabidopsis thaliana*, *Prunus persica*, *Vitis vinifera*, *Hordeum vulgare* and *Zea mays*). *Chlamydomonas reinhardtii* was used as an outgroup. Phylogenetic relationships were estimated using the Maximum Likelihood method implemented in PhyML+SMS with 1,000 bootstrap independent replicates. The seven defined clades are indicated, as well as potential subclades identified in this study (inner circles). Number on a node indicates the percentage of bootstrap when higher than 65% (black), or the posterior probabilities of major clades and subclades, according to a Bayesian phylogenetic inference performed on the same alignment (red) (**Supplementary Figure S1**). MLOs with one asterisk (*) have been experimentally demonstrated to be required for PM susceptibility (Büschges et al., 1997; Feechan et al., 2008; Wan et al., 2020), while MLOs with two asterisks (**) have been identified as main probable candidates for PM susceptibility (Pessina et al., 2016).

### 2.4 Transcriptional activity of *CsMLOs* in response to powdery mildew infection

#### 2.4.1 Sampling and RNA sequencing

An RNA-seq time series analysis of the infection of cannabis by PM was performed to characterize the transcriptional response of *CsMLO* genes. The experiment was conducted in a controlled environment at Organigram Inc, a Health Canada approved licensed producer (Moncton, New-Brunswick, Canada). Cannabis fan leaves from four-week-old vegetative plants (‘*Pineapple Express*’, drug-type I) were manually inoculated with *G. ambrosiae* emend. (including *G. spadiceus*) spores. Heavily infected leaves, loaded with fungal spores, were scraped against the surface of the leaves from the four-week-old plants to induce infection. Leaf samples (punch holes) were taken at five time points during the infection (n = 3 per time point): day zero (T0), six hours post-inoculation (6H), one day post-inoculation (1D), three days post-inoculation (3D) and eight days post-inoculation (8D). RNA samples were extracted from leaf tissues using the RNeasy® Plant Mini Kit (QIAGEN, Hilden, Germany) following the standard manufacturer’s protocol. Each mRNA extraction was treated with QIAGEN’s RNase-Free DNase Set (QIAGEN, Hilden, Germany), involving a first round of the TURBO DNA-free™ DNA Removal Kit through the extraction protocol and then two rounds of the DNA-free™ Kit (Life Technologies, Carlsbad, CA, USA). Fifteen individual RNAseq libraries were generated for the five time points sampled (n = 3 per time point). cDNA libraries were sequenced on a total of six lanes, using the Illumina HiSeq v4 technology (PE 125 bp) at the Centre d’expertise et de services Génome Québec (Montreal, QC, Canada). In total, ∼200 Gb of raw sequencing data were generated, which represents 1.728 billion of 2 x 125 bp paired-end sequences distributed across all 15 libraries (BioProject accession: PRJNA738505, SRA accessions: SRR14839036-50).

#### 2.4.2 Short-read alignment on the reference genome and differential gene expression

Raw sequencing reads were cleaned, trimmed, and aligned on the same reference genome as the one initially used to find our final *MLO* gene models (CBDRx) to estimate changes in transcript-specific levels of expression over the course of the infection. Specifically, mild trimming thresholds were applied to clean and trim all raw reads, using Trimmomatic v.0.34 (Bolger et al., 2014) with the following parameters: ILLUMINACLIP:$VECTORS:2:30:10, SLIDINGWINDOW:20:2, LEADING:2, TRAILING:2, MINLEN:60. The cleaned reads from our 15 individual libraries were aligned on the CBDRx reference genome using STAR v.2.7.6 (Dobin et al., 2013) in genome mode with default parameters and the official NCBI Cannabis annotation release 100 associated with the genome. Genome-wide raw read counts were obtained for each library using htseq-count v.0.11.1 (Anders et al., 2015) with the ‘intersection-non-empty’ mode.

Downstream analyses of differential gene expression patterns were conducted using the R packages ‘limma-voom’ (Law et al., 2014) and edgeR (Robinson et al., 2010) in RStudio v.1.3.1073 (RStudio Team, 2020). Reference sequences with insufficient sequencing depth were filtered out by keeping only the ones with more than five Counts Per Million (CPM) in at least three samples. This mild CPM threshold allowed the filtering of very low coverage genes without losing too much information in the dataset. This filtered dataset was normalized using the Trimmed Mean of M-values (TMM) method implemented in edgeR. A transformation of the data was then performed using the ‘limma-voom’ function, which estimates the mean-variance relationship for each transcript, allowing for better and more robust comparisons of gene expression patterns across RNA-seq libraries (Law et al., 2014). Each transcript was finally fitted to an independent linear model with log2(CPM) values as the response variable and the time point (zero, six hours, three days, and eight days post-infection) as the explanatory variable. All linear models were treated with limma’s empirical Bayes analysis pipeline (Law et al., 2014). Differentially expressed genes were chosen based on a False Discovery Rate (FDR, Benjamini–Hochberg procedure) < 0.01. Genomic regions corresponding to the curated CBDRx *MLO* gene models were extracted from the output of edgeR/limma-voom for each comparison made (i.e. each time point compared to T0) and looked for significant gene expression differences among *MLO* genes (FDR < 0.01). These expression differences in *MLOs* over the course of the infection were visualized in RStudio v.1.3.1073 (RStudio Team, 2020) on a scatter plot using CPM values (**Figure 4, Supplementary Figure S2**).

**Figure 4.**
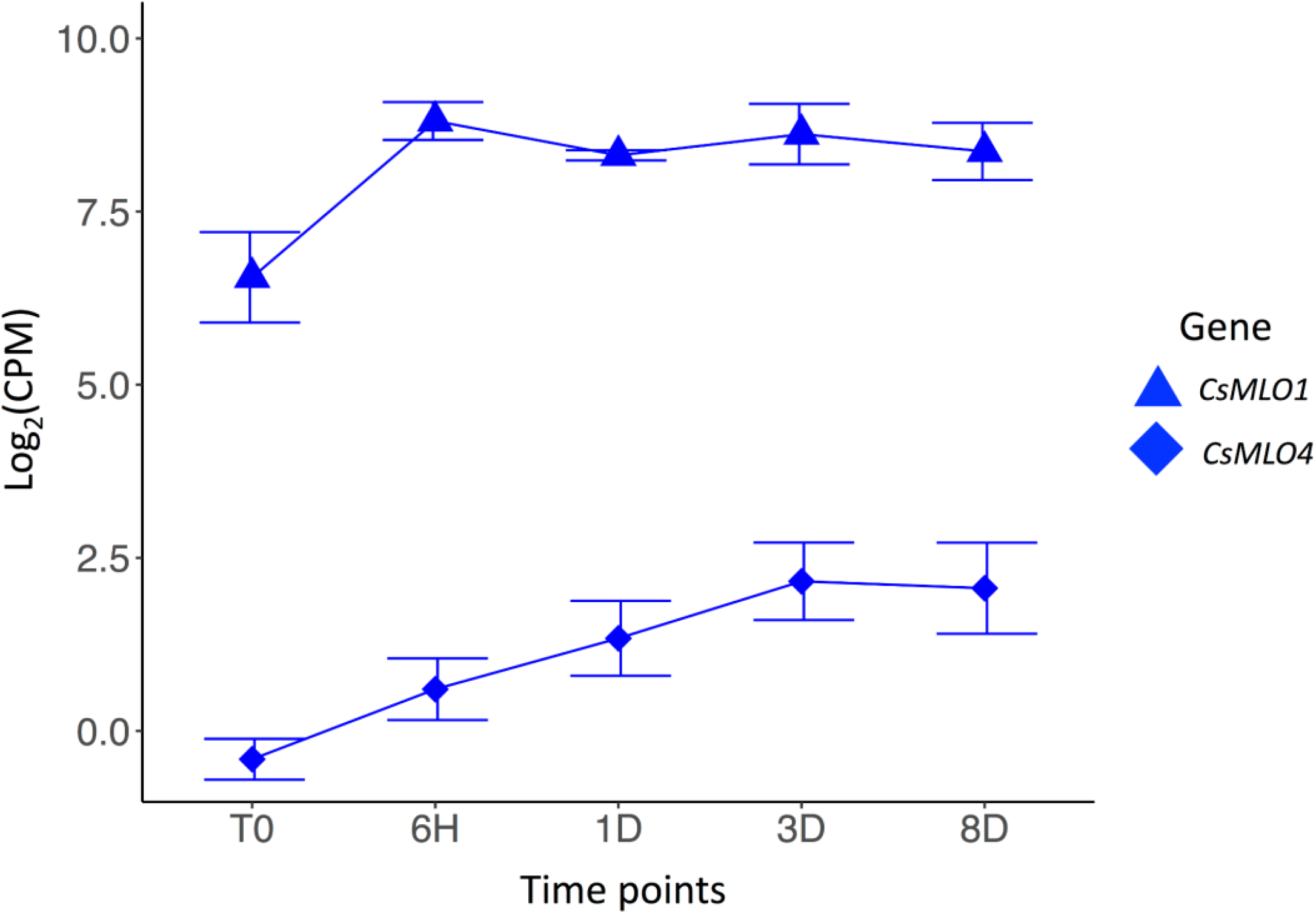
Transcriptomic response of clade V *CsMLO* genes following inoculations with powdery mildew. Time series analysis of the infection of *Cannabis sativa* leaves by PM, showing average gene expression for *CsMLO1* (blue triangles) and *CsMLO4* (blue diamonds). Gene expression is displayed on the y-axis as the average logarithmic value of the Counts Per Million (log2(CPM)) at each time point (displayed on the x-axis, n = 3 per time point). Time points: no infection / control (T0), 6 hours post-inoculation (6H), 24 hours post-inoculation (1D), 3 days post-inoculation (3D), and 8 days post-inoculation (8D).

### 2.5 Cloning of clade V *CsMLOs* for transient expression in *Nicotiana benthamiana* and confocal microscopy

The two selected *MLO* gene sequences (*CsMLO1* and *CsMLO4*) were synthesized commercially into the Gateway-compatible vector pDONR™/Zeo (Invitrogen, Carlsbad, CA, USA) (Hartley et al., 2000). The entry vectors were inserted into *Escherichia coli* OneShot® TOP10 cells (Invitrogen, Carlsbad, CA, USA) by chemical transformation according to the manufacturer’s instructions. Positive colonies were selected, and plasmid DNA was extracted with the EZ-10 Spin Column Plasmid DNA Miniprep Kit (Bio Basic Inc., Markham, ON, Canada). The extracted entry vectors were confirmed by PCR with the primers M13-F (5’-GTAAAACGACGGCCAGT-3’) and M13-R (5’-CAGGAAACAGCTATGAC-3’) as well as M13-F and MLO1_1-R (5’-ATGTGCCATTATAAATCCATGCCT-3’, this study).

The Gateway-compatible destination vector chosen was pB7FWG2.0, which is under the regulation of the 35S Cauliflower Mosaic Virus (CaMV) promoter and harbors the plant selectable marker gene *bar* (bialaphos acetyltransferase), which confer resistance against glufosinate ammonium. It also possesses a streptomycin and spectinomycin resistance gene for plasmid selection, and an EGFP-fusion in C-terminal, for visualization by confocal microscopy (Karimi et al., 2002). According to the manufacturer’s instructions, the destination vector was inserted into *Escherichia coli* One Shot® ccdB Survival™ cells (Life Technologies) by chemical transformation. Positive colonies were selected, and plasmid DNA was extracted with the EZ-10 Spin Column Plasmid DNA Minipreps Kit (Bio Basic Inc., Markham, ON, Canada). The extracted destination vectors were confirmed by PCR with the primers T-35S-F (5’-AGGGTTTCTTATATGCTCAACACATG-3’, Debode et al., 2013) and EGFP-C (5’-CATGGTCCTGCTGGAGTTCGTG-3’).

The two synthesized MLO gene sequences were then inserted into the vector pB7FWG2.0 through an LR clonase reaction following the manufacturer’s instructions (Invitrogen, Carlsbad, CA, USA). Plasmids were then transferred to *E. coli* OneShot® TOP10 cells (Invitrogen, Carlsbad, CA, USA) by chemical transformation according to the manufacturer’s instructions. Positive colonies were selected, and plasmid DNA was extracted with the EZ-10 Spin Column Plasmid DNA Minipreps Kit (Bio Basic Inc., Markham, ON, Canada). The extracted expression vectors were amplified by PCR with the primers 35S Promoter (5’-CTATCCTTCGCAAGACCCTTC-3’) and MLO1_1-R as well as MLO2_3-F (5’-TCTTTCAGAATGCATTTCAACTTGC-3’, this study) and EGFP-N (5’-CGTCGCCGTCCAGCTCGACCAG-3’). Sequencing of the PCR products above was performed to confirm the junction between the plasmid and the inserted gene and the junction between the inserted gene and the GFP, respectively.

The recombinant vectors were then transferred to *Agrobacterium tumefaciens* ElectroMAX™ LBA4404 cells by electroporation (Invitrogen, Carlsbad, CA, USA). The expression vector in *A. tumefaciens* was confirmed by PCR using the primers 35S Promoter and MLO1_1R and MLO2_3-F + EGFP-N. Cultures of transformed *A. tumefaciens* (*CsMLO1* and *CsMLO4*) were incubated with agitation at 28°C in LB broth containing spectinomycin (100 mg/mL) for 24 hours. The cultures were then centrifuged at 5000 x *g* for 5 minutes, the supernatant was discarded, and the pellet was resuspended in MgCl2 (10 mM). The cells were brought to an OD600 of 0.5 and incubated at room temperature for 2 hours with acetosyringone (200 µM). *Nicotiana benthamiana* plants, about two weeks old, were watered a few hours before infiltration, and the bacterial suspensions were administered using sterile 1 mL syringes (without needles) on the abaxial surface of the leaves. The plants were then returned to growth chambers, and the observation of epidermal cells was performed three days after infiltration using a confocal laser scanning microscope. Leaves were observed under a Leica TCS SP8 confocal laser scanning microscopy (Leica Microsystems). Images were observed through an HC PL APO CS2 40X/1.40 oil immersion objective at excitation/emission wavelengths of 488/503-521 nm.

### 2.6 *In silico* screening of clade V *CsMLO* sequence variants in 32 cannabis cultivars

Thirty-one distinct drug-type *Cannabis sativa* cultivars from Organigram Inc. (Moncton, New-Brunswick, Canada) and one industrial hemp variety (‘Anka’, UniSeeds, obtained from Céréla, Saint-Hughes, Québec, Canada) were screened to identify potential polymorphisms in clade V *CsMLOs* that could be associated with increased susceptibility or resistance to PM. Raw sequencing files (Illumina paired-end 125 bp) from these 32 cultivars (BioProject accession: PRJNA738519, SRA accessions: SRR14857079-110) were aligned on the reference CBDRx genome using ‘speedseq’ v.0.1.2 (Chiang et al., 2015). ‘bcftools view’ v.1.10.2 (Li, 2011) was used on the raw BAM alignment files to extract the genomic regions corresponding to the two clade V *CsMLO* genes, based on the positions of our manually curated CBDRx *MLO* gene models. Genotypes in these two *CsMLOs* were called for Single Nucleotide Polymorphisms (SNPs) and small insertions and deletions (INDELs) across the 32 cannabis cultivars using ‘bcftools mpileup’ v.1.10.2 (Li, 2011). Variants with a mapping quality < 20 and/or with a read depth > 500X were filtered out, and allelic frequencies for each variant were extracted using an in-house Python script (v.3.7.3, Python Software Foundation, 2020). Next, SNPGenie v.1.0 (Nelson et al., 2015) was used to estimate the ratio of non-synonymous to synonymous polymorphisms in the two clade V *CsMLO* genes across all 32 cultivars. Based on the output from SNPGenie, genomic positions of all non-synonymous polymorphisms found in the two genes were extracted using an in-house Python script (v.3.7.3, Python Software Foundation, 2020). Visual representations of these polymorphisms at the gDNA and amino acid levels were prepared using Microsoft © PowerPoint v.16.49 (Microsoft Corporation, Redmond, Washington, U.S.).

## 3 Results

### 3.1 *CsMLO* gene identification and genomic localization

Through careful manual curation, we were able to identify a total of 14, 17, 19, 18 and 18 distinct *CsMLO* genes in the genomes of CBDRx, Jamaican Lion female, Jamaican Lion male, Purple Kush, and Finola, respectively. Our final manually curated *CsMLO* genes were numbered 1-15 based on chromosomal positions in CBDRx, from chromosome 1 through chromosome X (**Table 1**, **Supplementary Tables S1-S5, Supplementary Files S1-S3**). *CsMLO* gene numbers and IDs in all four other genomes (Finola, Jamaican Lion female, Jamaican Lion male, Purple Kush) were based on homology relationships, supported by our phylogenetic and orthology analyses (see Materials and Methods for details). We also physically located each set of *CsMLO* genes on the chromosomes of CBDRx, Finola and Purple Kush (**Figure 1**). Our results show that eight of the 10 chromosomes harbor evenly-spaced *CsMLO* genes, chromosomes 5 and 9 being the only ones not carrying MLO genes. We were able to anchor all 14 CBDRx *CsMLO* genes on their respective chromosomes, while two (*CsMLO12-FN*, *CsMLO15-FN*) and six (*CsMLO1-PK_B*, *CsMLO2-PK*, *CsMLO4-PK*, *CsMLO6-PK*, *CsMLO7-PK*, *CsMLO12-PK*) *MLO* genes were located on unanchored contigs in Finola and Purple Kush, respectively (**Figure S1, Table 1, Supplementary Tables S1, S4 and S5**). The CBDRx genome showed an absence of MLO15, while this gene was present in all other cannabis genomes. We identified homology and paralogy relationships among all *CsMLO* sequences, which revealed potential duplication patterns in certain genomes. Overall, we detected paralogs for *CsMLO1*, *CsMLO5*, *CsMLO9*, *CsMLO10*, *CsMLO12* and *CsMLO13* in the genomes of Finola, Purple Kush, Jamaican Lion male and Jamaican Lion female (*CsMLO* paralogs were designated with A/B suffixes, see **Table 1, Supplementary Tables S1-S5**). CBDRx remained the only genome in which we did not identify *CsMLO* paralogs. Most of the *CsMLO* genes had syntenic positions across all three chromosome-level genome assemblies, with the exception of *CsMLO9*, *CsMLO10* and *CsMLO14* (**Figure 1**). *CsMLO14* was the only *CsMLO* gene identified across all five genomes that had three different locations in the three genomes, i.e. chromosome X in CBDRx, chromosome 1 in Finola and chromosome 4 in Purple Kush. Partial/incomplete genes were also noted in the genomes of Finola (*CsMLO3-FN-B* and *CsMLO10-FN-A*) and Purple Kush (*CsMLO1-PK-B*, *CsMLO2-PK* and *CsMLO5-PK-A*).

### 3.2 *CsMLO* gene structure and protein characterization

Manually curated *CsMLO* genes identified in CBDRx ranged in size between 3,567 bp (*CsMLO14-CBDRx*) and 49,009 bp (*CsMLO5-CBDRx*), with an average size of 11,240 bp and a median size of 5,431 bp (**Table 1, Supplementary Tables S1-S5**). The structural organization of these *CsMLO* genes is depicted in **Figure 2**. The number of exons varied between 12 and 15, with some of the exons showing signs of fusion in certain genes (**Figure 2**). The number of amino acid residues in these *CsMLO* gene sequences varied between 509 and 632. Intron size varied considerably, with 11 introns belonging to four different *CsMLO* genes (*CsMLO4*, *CsMLO5*, *CsMLO6*, *CsMLO10*) exhibiting a length greater than 1,000 bp. *CsMLO5* had the longest introns: intron 5 (15.6 kb), intron 9 (7.9 kb), intron 12 (15.9 kb), and intron 14 (6.3 kb) (**Figure 2**). The *CsMLO* genes that we characterized in the four other genomes (Finola, Jamaican Lion female and male, Purple Kush) were consistently similar to the ones identified in CBDRx in terms of length, intron and exon structural organization and genomic localization (**Supplementary Files S1-S3**). The longest *CsMLO* gene characterized among all our manually curated gene models belonged to Jamaican Lion (female), with a total length of 49,673 nucleotides (*CsMLO5-JL*).

Proteins encoded by all identified *CsMLO* genes comprised seven transmembrane domains (TMs) of similar lengths (**Table 1, Supplementary Tables S1-S5**). The only exceptions were found in the genomes of Finola and Purple Kush in which we identified a total of four partial *CsMLO* genes, which harbored less than seven TMs: CsMLO3-FN-B (5 TMs), CsMLO10-FN-A (3 TMs), CsMLO1-PK-B (4 TMs), CsMLO5-PK-A (6 TMs). Similarly, all identified *CsMLO* proteins were predicted to be localized in the cell membrane by several online prediction servers, such as Plant-mPLoc (Chou and Shen, 2010), DeepLoc 1.0 (Almagro Armenteros et al., 2017), and YLoc-HighRes (Briesemeister et al., 2010) (**Table 1, Supplementary Tables S1-S5**). Three partial/incomplete sequences were also predicted to localize elsewhere, such as the chloroplasts and nucleus by Plant-mPLoc for CsMLO1-PK-B, and the endoplasmic reticulum by DeepLoc 1.0 for CsMLO5-PK-A and CsMLO3-FN-B. We assumed that these three predictions were unreliable as they were made using incomplete sequences and not present unanimously throughout all prediction servers. No signal peptide (SignalP 5.0, Almagro Armenteros et al., 2019a), mitochondrial transit peptide, chloroplast transit peptide, or thylakoid luminal transit peptide (TargetP 2.0, Almagro Armenteros et al., 2019b) were predicted in any of the CsMLO protein sequences (results not shown). The invariable 30 amino acid residues previously described in Elliott et al., 2005 were identified in all CsMLOs (**Table 1, Supplementary Tables S1-S5**). Across all identified CsMLOs (excluding the five partial sequences), the amount of conserved amino acids varied between 25 and 30, and 74.1% (60/81) of CsMLO sequences possessed all 30 amino acids. In Kusch et al., 2016, a larger dataset of MLO proteins was analyzed and thus identified 58 highly conserved amino acids, rather than invariant, showing that substitutions are possible. These 58 amino acid residues were also screened in all CsMLOs (**Table 1, Supplementary Tables S1-S5**). Across all identified CsMLOs (excluding the five partial sequences), the amount of conserved amino acids varied between 52 and 58, and only 23.5% (19/81) of CsMLO sequences possessed all 58 amino acids. However, 74.1% (60/81) possessed 57 or more of the conserved amino acids.

### 3.3 Phylogenetic analysis of CsMLOs

We performed a phylogenetic analysis on the curated cannabis MLO proteins identified among the five genomes (CsMLOs). The dataset was completed with the MLO protein family from *Arabidopsis thaliana* (AtMLO, Devoto et al., 2003), *Prunus persica* (PpMLO, Pessina et al., 2014), *Vitis vinifera* (VvMLO, Feechan et al., 2008), *Hordeum vulgare* and *Zea mays* (HvMlo and ZmMlo, Kusch et al., 2016), using the *Chlamydomonas reinhardtii* MLO as the outgroup. Phylogenetic tree construction was performed using the PhyML+SMS algorithm, which confirmed the seven known clades of MLO proteins (**Figure 3**), with bootstrap values equal or greater than 97% (except for clade I, supported with a bootstrap value of 77%). Clade numbers from I to VII were assigned according to the previous study of Kusch et al. (2016). Previous studies have reported the presence of an eighth clade (e.g., Pessina et al., 2014), clustering with clade VII in other papers. Here, we followed the seven clades nomenclature, but this potential eighth clade would correspond to one of the two clade VII subclades. Indeed, potential subclades were also identified in our study, indicated as separate lines in the inner circle of **Figure 3**, which were also supported with high values of bootstrap (equal or greater than 83%, with the exception of one clade V subclade, supported with a value of 67%). The same clades and subclades were also found to be supported by high posterior values equal or greater than 77% (indicated in red), following phylogenetic tree construction using MrBayes (also see **Supplementary Figure S1**). Apart from one clade II subclade that appeared monocot-specific, all subclades depicted here included CsMLO, as well as PpMLO and VvMLO. In each subclade, all CsMLOs clustered together with bootstrap values of 100. Two subclades were found in the phylogenetic clade V, containing all the dicot MLO proteins experimentally shown to be required for PM susceptibility (Acevedo-Garcia et al., 2014). In one of those clade V subclades, three of the cannabis genomes were found to harbor two near-identical genes. All cannabis genomes except CBDRx were found to include one *MLO* gene grouping with clade IV, which contains all monocot MLO proteins acting as PM susceptibility factors (**Figure 3**).

### 3.4 Transcriptional reprogramming of cannabis *MLOs* in response to *Golovinomyces ambrosiae* infection

We conducted an RNA-seq analysis to look at the transcript abundance of *CsMLO* genes in leaves of the susceptible cannabis cultivar “Pineapple Express” during infection by *G. ambrosiae* emend. (including *G. spadiceus*). Five time points were investigated, corresponding to key stages of the infection: 0 (control), 6 hours post-inoculation (hpi) (conidia germination and appressoria formation), 24 hpi (haustoria formation), 3 day post-inoculation (dpi) (secondary hyphae formation), and 8 dpi (secondary haustoria, secondary appressoria and conidiophore/conidia formation). No significant expression was detected at any time point for *CsMLO6*, *CsMLO8*, *CsMLO11*, and *CsMLO15* (in this particular case, reads were aligned to the genome of Jamaican Lion male, as the CBDRx genome is devoid of *CsMLO15*). While *CsMLO3* and *CsMLO11* were expressed, no up- or down-regulation was observed under our conditions. Nine genes, namely *CsMLO1*, *CsMLO2*, *CsMLO4*, *CsMLO5*, *CsMLO7* and *CsMLO9* were found to be significantly differentially expressed (FDR < 0.05) after inoculation with the pathogen (**Figure 4, Supplementary Figure S2**). In the case of clade V genes, *CsMLO1* showed a peak of 2.23-fold up-regulation at 6 hpi (FDR = 2.83×10^-4^), and remained somewhat constant for the remaining of the infection, while the expression of *CsMLO4* increased steadily between T0 and 3 dpi, reaching a peak of 2.02-fold up-regulation (when compared to T0) and remained constant at 8 dpi (FDR = 7.87×10^-4^).

An analysis of the 2 kb upstream region of all *CsMLO* genes identified through the five cannabis genomes revealed the presence of key regulatory motifs with functions related to environmental/hormonal response (e.g., ABRE, AuxRR-core, ERE, GARE, P-box, TATC-box, TGA-element, GT1-motif, G-box, light response elements), stress and defense response (TC-rich repeats, MBS, ARE, GC-motif, LTR element), developmental regulation (circadian, HD-Zip, CCAAT-box, MSA-like), seed-specific metabolism (O2-site, RY-element) and wound response (WUN-motif). In total, 82 (95%) *CsMLO* genes had at least one motif related to environmental/hormonal response, while 80 (93%) *CsMLO* genes had a MYB-related sequence, a motif typically involved in development, metabolism and responses to biotic and abiotic stresses (Dubos et al., 2010) (**Supplementary Table S6**). We found the presence of at least one *cis*-acting element involved in gene overexpression by biotic and abiotic factors (ABRE, CGTCA, TGACG, TCA) in 100% of the 13 *CsMLO* genes from clade V identified in all five genomes. The analysis of protein domains, their location in the protein and the overall topology of each gene ultimately revealed a consistent pattern among all *CsMLO* genes.

### 3.5 Subcellular localization of clade V *CsMLOs*

The subcellular localization of clade V MLOs, such as CsMLO1 and CsMLO4, was first analyzed using online tools. As mentioned previously, these analyses predicted that CsMLO1 and CsMLO4 possessed seven transmembrane domains and were localized in the plasma membrane (**Table 1, Supplementary Tables S1-S5**). To determine the subcellular localization of CsMLO1 and CsMLO4 *in planta*, we constructed two vectors under the control of the CaMV 35S promoter where the coding sequences of CsMLO1 and CsMLO4 were fused to enhanced green fluorescent protein (EGFP) in C-terminal (*35S::CsMLO1-EGFP* and *35S::CsMLO4-EGFP*). The agroinfiltration-based transient gene expression system was used to transform *Nicotiana benthamiana* leaves with each construct. The epidermal cells were observed three days after transformation for GFP signal using confocal laser scanning microscopy. The CsMLO1-EGFP fusion protein was observed in the cell periphery as well as throughout the cell forming networks in a punctuate pattern, while the CsMLO4-EGFP fusion protein was observed solely in the cell periphery in a defined way (**Figure 5**).

**Figure 5.**
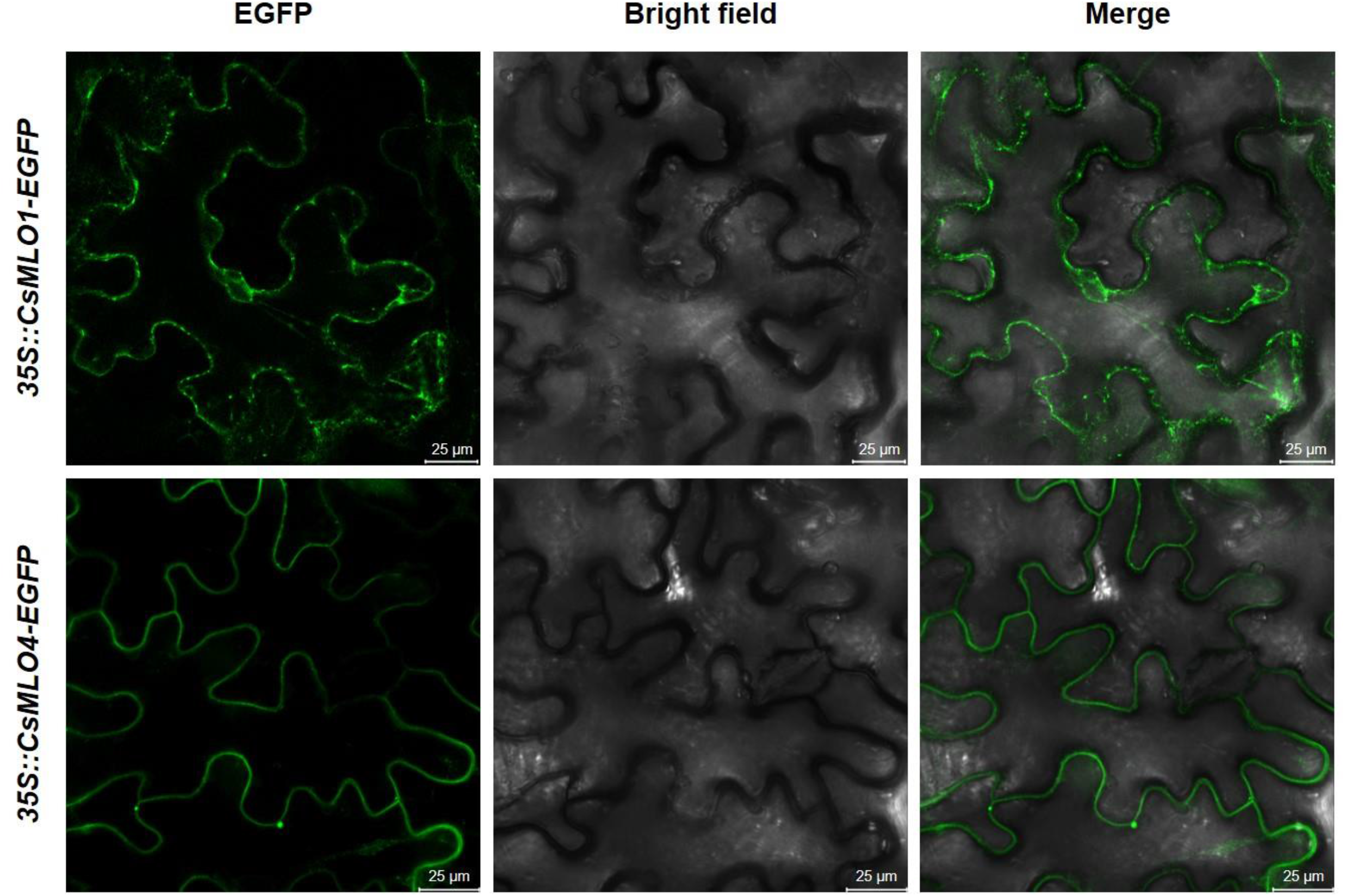
Subcellular localization of *CsMLO1* and *CsMLO4* as observed by confocal laser scanning microscopy. Transient expression of *35S::CsMLO1-EGFP* and *35S::CsMLO4-EGFP* constructs in *Nicotiana benthamiana* leaf epidermal cells shown 3 days after agro-infiltration. Scale bars = 25 μm.

### 3.6 Polymorphism analysis of clade V *CsMLOs* among 32 cultivars

Comparison of 32 distinct cannabis cultivars to the CBDRx reference genome to detect polymorphisms in *CsMLO* clade V genes revealed a total of 337 (*CsMLO1*, 101 indels and 236 SNPs) and 852 (*CsMLO4*, 154 indels and 698 SNPs) polymorphisms, mainly located in the last cytosolic loop of the protein, near the C-terminus. Among these polymorphic loci, we identified 14 and 12 non-synonymous SNPs for *CsMLO1* and *CsMLO4* respectively (**Supplementary Table S7**). The only SNP that was not located near the C- or N-terminus was found in *CsMLO1*, in the second cytoplasmic loop. All other SNPs in *CsMLO1* were distributed in the first cytoplasmic loop (three non-synonymous SNPs), the second extracellular loop (four non-synonymous SNPs) and the last cytoplasmic loop, near the C-terminus (six non-synonymous SNPs, **Figure 6**). The scenario for *CsMLO4* is slightly different, with a single non-synonymous SNP identified in the second extracellular loop and the remaining 11 SNPs all located in the last cytoplasmic loop, near the C-terminus (**Figure 6**). About half of these non-synonymous SNPs in both genes induced a change in amino acid charges or polarity, with seven (50%) and five (42%) SNPs having a change in electric charges in *CsMLO1* and *CsMLO4*, respectively. None of the SNPs identified in either of the two *CsMLO* clade V genes had an impact on conserved amino acids in the proteins (Elliott et al., 2005; Kusch et al., 2016). Overall, allele frequencies associated with these SNPs in *CsMLO1* and *CsMLO4* showed an even distribution of reference and alternate alleles throughout the 32 cultivars. There was one exception with ‘Ultra Sour’, which was identified as the only homozygous cultivar for the alternate allele in three non-synonymous SNPs found in exons 1 (G40E) and 3 (P96Q, P111T).

**Figure 6.**
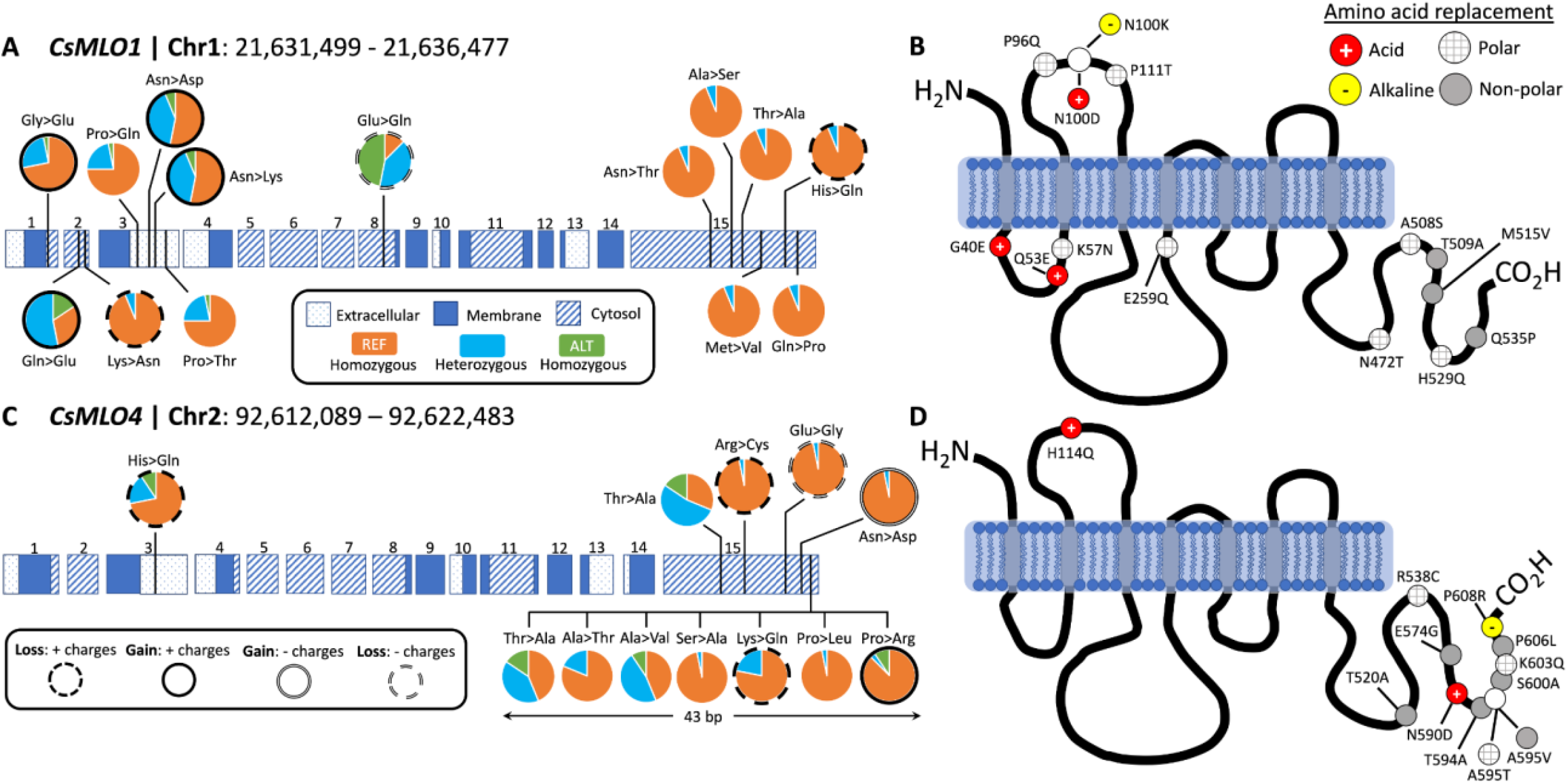
Protein structure and polymorphism analysis of the two clade V genes (*CsMLO1* and *CsMLO4*) in 32 distinct cannabis cultivars aligned against the CBDRx reference genome. Panels A and B exhibit all non-synonymous nucleic acid substitutions (A) and amino acid replacements (B) identified in *CsMLO1* (located on chromosome 1) from 32 hemp and drug-type cannabis cultivars. Panels C and D exhibit all non-synonymous nucleic acid substitutions (C) and amino acid replacements (D) identified in *CsMLO4* (located on chromosome 2) from the same 32 hemp and drug-type cannabis cultivars as shown on panels A and B. On panels A and C (gDNA), pie charts display, for each SNP along the coding sequence, the genotype frequencies calculated among the 32 cultivars. Colored horizontal rectangles on panels A and C represent the exons, numbered from 1 to 15: dotted rectangles represent extracellular domains of the resulting protein, while blue and striped rectangles represent membrane and cytoplasmic domains, respectively. Genotype frequency color code on panels A and C: samples called as homozygous for the reference allele are depicted in orange, samples called as heterozygous are depicted in light blue, and samples called as homozygous for the alternate allele are depicted in green. On panels B and D (amino acid sequence), the gray vertical rectangles depict transmembrane domains and the solid black curves depict cytoplasmic and extracellular loops. Resulting amino acid replacements on panels B and D are color-coded according to the gain or loss of polarity: red circles with white cross display a replacement with an acidic amino acid, yellow circles with black hyphen represent a replacement with an alkaline amino acid, gray circles display a replacement with a non polar amino acid, and gray-striped white circles represent a replacement with a polar amino-acid.

## 4 Discussion

Considering the role of specific *MLO* genes in flowering plants’ susceptibility to powdery mildew, one of the most prevalent pathogens in indoor cannabis productions (Punja et al., 2019), our primary goal was to structurally and functionally characterize this gene family at a manual-curation level in multiple cannabis genomes. In order to develop mitigation strategies aimed at reducing the deleterious impacts of the pathogen on cannabis productions, and in an attempt to better understand other functional roles of *CsMLOs*, the first step consisted in identifying the exact number and structure of these genes in different genetic backgrounds. Our results first showed that *CsMLO* numbers are variable across different cannabis types. Second, they showed that two distinct clade V genes were present in all genomes (with paralogs in certain cultivars) and that these clade V genes possessed *cis*-acting elements typically overexpressed by biotic and abiotic factors. These specific elements in the promoter regions of clade V *CsMLOs* allow them to be responsive to experimental PM infection, which we validated in the context of an infection time point experiment.

### 4.1 *CsMLOs* and the importance of manual curation

We used 15 *Arabidopsis thaliana* MLO protein sequences to mine the genomes of five cannabis cultivars of four different types (THC-dominant, balanced THC:CBD, CBD-dominant, food-oil hemp), yielding a sum of 86 *CsMLOs* across all genomes (**Table 1, Supplementary Tables S1-S5**). According to our data, these genes are organized into 15 *CsMLO* homologs in total (**Figures 1 and 3, Supplementary Tables S1-S5**). We were not able to retrieve *CsMLO15* in CBDRx, making this genome devoid of clade IV *MLO*. Whether this is an artifactual gene loss resulting from the genome assembly and cleaning process or a biological reality in this specific cultivar remains to be verified with additional sequencing data. If this is a biological reality, it could suggest that these genes potentially have redundant and similar functions or are imbricated into functional networks buffered by redundancy (Tully et al., 2014; AbuQamar et al., 2017). The loss of a gene in the genome of CBDRx in this context could potentially be phenotypically less detrimental. Gene length also varied with a 14-fold difference between the shortest and longest sequence, with two genes (*CsMLO5* and *CsMLO10*) exhibiting multiple unusually long introns (> 10,000 nucleotides, **Figure 2**). Intron length distribution across the 15 *CsMLOs* showed considerable variability, which explained the significant variations observed in gene length across all *CsMLOs*. Plant introns are typically relatively short and rarely extend beyond 1 kb, making these large *CsMLO* introns up to 10 times longer than typical plant exons (Wu et al., 2013). However, the numbers and positions of exons and introns for the same homologs across the five genomes were highly conserved. We found that cannabis genes tend to have on average five introns (median number of introns across the genome = 4), indicating that *CsMLOs* have three times the number of introns found in a typical gene from the cannabis genome. This could indicate that those introns are likely to play an important functional role and, thus, may be a significant aspect of gene regulation (Seoighe and Korir, 2011). On the other hand, selection against intron size could be counterbalanced in *CsMLOs* by a selective preference for larger introns which correlates with more regulatory elements and a more complex transcriptional control (e.g., in *Vitis vinifera*, Jiang and Goertzen, 2011). Even though the complete *CsMLO* gene catalog could be retrieved in each genome through gene prediction algorithms combined with targeted BLAST searches, the precise characterization of each gene structure (i.e. start/stop codons, coding sequence, exon-intron boundaries) could not be achieved without multiple efforts of manual curation.

Automated gene structure prediction algorithms are often considered sufficiently reliable to recover the complete genome-wide repertoire of genes of a given sample. However, as repeat content, size, and structural complexity of those genes increase (i.e. numerous small exons delimited by long introns, as observed in *CsMLOs*), errors are increasingly likely to occur and thus impair the accuracy of automated annotations (Guigó et al., 2000; Pilkington et al., 2018). Because 13% of the *CsMLOs* identified here had introns larger than 10 kb (*CsMLO5* and *CsMLO10*), and two additional genes (*CsMLO4* and *CsMLO6*) had introns larger than 1,000 bp, most of these *CsMLOs* were mispredicted by automated gene prediction algorithms (**Supplementary Tables S1, S2 and S3**). These algorithms typically have 10 kb as the default maximum intron length (e.g., in MAKER2, Holt and Yandell, 2011). In comparison, *Arabidopsis thaliana* genome annotation version 10 indicates that there are 127,854 introns in the nuclear genes, and of these, 99.23% are less than 1,000 bp, while only 16 introns are larger than 5 kb (NCBI accession: GCF_000001735.4, TAIR10.1). Long multi-exon genes having long introns end up fragmented into several shorter “genes” by these programs, thus inflating the actual number of genes within the family. The severity of such errors is influenced by various factors such as the quality of the assembly (Yandell and Ence, 2012; Treangen and Salzberg, 2012) and the availability and quality of extrinsic evidence (e.g., RNA-seq, orthologous sequences). While assembly quality is influenced by genome size and repeat content (Tørresen et al., 2019; Whibley et al., 2021), the disparity in the number of mispredicted genes observed in this study is also likely to be related to differences in sequencing technology (Oxford Nanopore vs. Pacific Biosciences), sequencing depth, algorithms used for *de novo* assembly, and simple base calling accuracy. Overall, these results revealed that cannabis possesses an extensive repertoire of *MLOs* characterized by significant gene size variations across all family members. These results also demonstrate the importance of manual curation when working with automatically generated gene models. Identifying the complete and detailed set of *CsMLOs* for each genome allowed the possibility to assess synteny and evolutionary patterns among *Cannabis sativa* and other plant species.

### 4.2 Evolutionary dynamics of *CsMLOs*, gene duplications and potential implications

The vast majority of studies investigating phylogenetic relationships within the *MLO* gene family usually classifies its members in seven clades. A few times, an extra clade has been proposed, or various subclades have been identified, but no consensus has been reached yet. In our study, we classified genes according to the seven clades, and identified some potential subclades presented in **Figure 3**, most of which appear concordant with results from other studies. In two previous studies (Polanco et al., 2018; Rispail and Rubiales, 2016), clade I is divided in two subclades (Ia and Ib). Subclade Ia would correspond in our tree to the two subclades respectively including *CsMLO12* and *CsMLO9*, while subclade Ib would correspond to the subclade that includes *CsMLO2*. However, Iovieno et al. (2016) divided clade I in three subclades (Ia, Ib and Ic). Subclade Ia from Iovieno et al. (2016) appears to correspond to subclade Ib from Polanco et al. (2018), and in our tree is represented by the subclade including *CsMLO2*. Subclades Ib and Ic from Iovieno et al. (2016) would correspond to subclade Ia from Polanco et al. (2018) and would be represented by the subclades including *CsMLO12* and *CsMLO9*, respectively. In Iovieno et al. (2016), clade II is divided into 15 subclades (named from IIa to IIq excluding IIj). Subclades IIa and IIb would be represented in our tree by the subclade including *CsMLO6*; subclades IIc, IId and IIe would be represented by the monocot-specific subclade; subclades IIf, IIg and IIh would be represented by the subclade including *CsMLO13*; and subclades IIi to IIq would be represented by the subclade including *CsMLO14*. Still in Iovieno et al. (2016), clade III is divided into 3 subclades (IIIa, IIIb and IIIc), subclades IIIb and IIIc corresponding in our tree to the subclades including *CsMLO8*, and *CsMLO7*, respectively. Subclade IIIa would correspond to the two immediate outlying monocot sequences (*ZmMlo2* and *ZmMlo3*), while our two next outlying monocot sequences (*HvMlo2* and *ZmMlo4*) would correspond to clade VIII (in Iovieno et al., 2016, as there is no consensus on clade VIII). Clade IV has also been divided into two subclades (IVa and IVb) which are grouped together in our tree, subclade IVa from their study simply corresponding to monocot sequences and subclade IVb corresponding to dicot sequences. Clade V has been divided into 3 subclades (Va, Vb and Vc), for which subclade Va corresponds in our tree to the subclade including *CsMLO1*, while subclade Vb would correspond to the two outlying sequences from the other subclade from our tree (*PpMLO3* and *VvMLO3*), and subclade Vc would correspond to the remaining of this subclade that includes *CsMLO4*. Clade VI is not divided into subclades in this paper, while we clearly identified two subclades in our tree, one including *CsMLO5* and one including *CsMLO3*. Clade VII has been divided into two subclades in Iovieno et al. (2016), and the same division can be found in our tree, with Iovieno’s subclade VIIa corresponding to the subclade that includes *CsMLO11*, while subclade VIIb would correspond to the subclade including *CsMLO10*. In Zheng et al. (2016), a different clade VIII had been defined, which would correspond to Iovieno et al. (2016) subclade VIIa, while Zheng et al. (2016) clade VII would correspond to Iovieno et al. (2016) subclade VIIb. In most studies, this eighth clade has been merged within clade VII, which is also the case here. As described above, a distinct clade VIII was also defined as a monocot-specific clade in Iovieno et al. (2016), making the use of an 8-clades system confusing. In our opinion, the clade VIII described by Iovieno et al. (2016) could be considered as a subclade of clade III, represented in our tree by the most “diverged” sequences in this clade, *HvMlo2* and *ZmMlo4*.

Manual curation of *CsMLOs* across the five studied genomes revealed with exactitude their respective genomic localization, showing an overall conserved syntenic pattern, except for two genomes, Finola and Purple Kush, which exhibited distinct chromosome localizations for specific *CsMLO* orthologs, as compared to the rest of the genomes. The number of *CsMLOs* per genome, ranging from 14 to 18, was comparable to other plant genomes, such as *Arabidopsis thaliana* (15, Chen et al., 2006), *Vitis vinifera* (14, Feechan et al., 2008), *Cucumis sativus* (13, Zhou et al., 2013), *Solanum lycopersicum* (15, Chen et al., 2014), *Hordeum vulgare* (11, Kusch et al., 2016), *Medicago truncatula* (14, Rispail and Rubiales, 2016), *Cicer arietinum* (13, Rispail and Rubiales, 2016), *Lupinus angustifolius* (15, Rispail and Rubiales, 2016), *Arachis* spp. (14, Rispail and Rubiales, 2016), *Cajanus cajan* (20, Rispail and Rubiales, 2016), *Phaseolus vulgaris* (19, Rispail and Rubiales, 2016) and *Vigna radiata* (18, Rispail and Rubiales, 2016). The genomes of Finola and Purple Kush, however, exhibited certain anomalies. Firstly, we found a greater proportion of genes located on unanchored contigs in Finola (11%) and Purple Kush (33%) as compared to CBDRx (0%). Secondly, two paralog pairs, one in Finola (*CsMLO10-FN-A* and *CsMLO10-FN-B*) and one in Purple Kush (*CsMLO9-PK-A* and *CsMLO9-PK-B*), had one of the two paralogs located on a different chromosome (**Figure 1**).

The genome-wide distribution of *CsMLOs* described here did not suggest the involvement of tandem duplications as a predominant mechanism of emergence for *MLOs* in cannabis, as suggested in other taxa (e.g., Liu and Zhu, 2008; Pessina et al., 2014; Rispail and Rubiales, 2016). Indeed, recent bioinformatic analyses suggested that segmental and tandem duplications were a widespread mechanism for the expansion of the *MLO* gene family in diverse plant species, spanning from algae to dicots (Shi et al., 2020). For example, clear evidence of tandem duplication events have been detected in *P. vulgaris* and *V. radiata*, and in *M. domestica*, respectively (Rispail and Rubiales, 2016; Pessina et al., 2014). We did not find evidence that tandem duplications were widespread in *CsMLOs*, as most of the genes were evenly spread out in the genomes, with multiple other unrelated genes in between these *CsMLOs*. Some of the *CsMLOs* were located physically close to one another and they were genetically related, but not similar enough (< 85%) to be considered as tandem duplicates. In this case, segmental duplication appears to be a more likely mechanism of emergence for *CsMLOs*, although we did not specifically search for segmental duplications in the present study. In total, four (4.7%) *CsMLOs* across the five genomes were located in tandem duplications (**Figure 1**). These four putative tandem gene duplications were located in Finola only, on chromosome X (*CsMLO13-FN-A* and *CsMLO13-FN-B*) and on chromosome 2 (*CsMLO3-FN-A* and *CsMLO3-FN-B*). Other *CsMLOs* that could represent potential tandem gene duplications in Jamaican Lion male (*CsMLO1-JLm*, *CsMLO12-JLm, CsMLO13-JLm*) and female (*CsMLO1-JL*, *CsMLO10-JL*) all had the two paralogs located on different contigs that were, on average, longer than 2 Mb each. These different contigs containing two *CsMLO* paralogs typically harbored large sequences (> 10 kb) of high homology (> 95% similarity), which could suggest that they are either the result of segmental duplications, or that they represent two copies of highly polymorphic loci (Fan et al., 2008; Lallemand et al., 2020). We did not find evidence of tandem duplications in the genome of Purple Kush, indicating that this genome is potentially more fragmented than the others. The duplication of *CsMLO13* (clade II) on chromosome X is of potential interest as it was duplicated in the male genomes only (Finola and Jamaican Lion male, **Supplementary Tables S3 and S5**). *MLO* clade II genes originally evolved in ancient seed-producing plants, suggesting that genes from this clade could have sex-related functions (Feechan et al., 2008; Jiwan et al., 2013; Zhou et al., 2013). On the other hand, this male-specific duplication could represent a technical artefact caused by the fact that the sequence of *CsMLO13* on chromosome Y was concatenated with its homologous version on chromosome X, thus producing a false tandem duplication. To our knowledge, no studies have documented in detail the structure and evolutionary relationships between orthologous *MLOs* in a group of genomes from the same species. Plus, information on sex-specific distribution of *MLOs* in comparable heterogametic sex plant systems is scarce, which makes the interpretation of this finding difficult. This could ultimately indicate that clade II *MLOs* may not be solely related to seed production or development, as suggested in other systems (Kusch et al., 2016). Recently, a clade II *MLO* was shown to be dictating PM susceptibility in mungbean (Yundaeng et al., 2020). However, apart from this example, only clade V *MLOs* have been shown to be involved in this trait in dicots. In this study, one cannabis clade V *MLO*, *CsMLO1*, appears to be duplicated in three different genomes (three out of the four THC-producing cultivars). If not an assembly artefact, the presence of such an additional copy of a clade V *MLO* would make it tedious to obtain complete immunity to PM. In other plants such as *Arabidopsis*, inactivation of all clade V *MLOs* is required to achieve complete immunity, even though these genes unevenly contribute to susceptibility, with *AtMLO2* playing a major role (Consonni et al., 2006). The presence of a single copy of *CsMLO1* in the industrial hemp variety Finola and the CBD-dominant CBDRx (which ancestry has been suggested to be 11% hemp, Grassa et al., 2021) could suggest that hemp is de facto less susceptible or that attaining this target in hemp could be easier to achieve. On the other hand, both Jamaican Lion cultivars investigated here have been described as being highly resistant to PM (McKernan et al., 2020), even though both have this extra, apparently functional, *CsMLO1* copy. In this particular case, resistance to PM is likely due to other genetic factors than a loss-of-susceptibility that would have been obtained through deletion/mutation of clade V *MLOs* (see below).

### 4.3 Upregulation of clade V MLOs triggered by powdery mildew infection

The transcriptomic response of *Cannabis sativa* to PM performed here revealed that clade V *CsMLOs* are rapidly triggered, at least six hours post-inoculation, upon infection by the pathogen (**Figure 4**). Our results revealed that the two clade V *CsMLO* genes responded with different activation patterns, potentially suggesting specific roles. As described above, all three clade V *MLO* genes need to be inactivated in order to achieve complete immunity against PM in *Arabidopsis* (Consonni et al., 2006). However, in other plants, not all members of clade V are *S*-genes, and the inactivation of only a subset of the clade V *MLO* genes is required (Bai et al., 2008; Pavan et al., 2011). In those cases, the precise identification of the exact genes involved in susceptibility needs to rely on additional criteria. A shared element of all *MLO* genes involved in PM susceptibility is that they respond to fungal penetration, showing significant upregulation as soon as six hours after inoculation (Piffanelli et al., 2002; Bai et al., 2008; Zheng et al., 2013), and candidate genes can thus be identified based on increased expression (Feechan et al., 2008; Pessina et al., 2014). Indeed, being an *S*-gene, the expression of *MLO* is necessary for the successful invasion of PM (Freialdenhoven et al., 1996; Lyngkjær et al., 2000; Zellerhoff et al., 2010). For instance, in watermelon, upregulation was only observed for one clade V *MLO* (out of five), *ClMLO12*, and only at time points corresponding to nine and 24 hours after inoculation with *Podosphaera xanthii*, making it the prime candidate dictating PM susceptibility (Iovieno et al., 2015). In apple, three genes (including two out of the four clade V *MLOs*, *MdMLO11* and *MdMLO19*) were found to be significantly up-regulated after inoculation with PM, reaching about 2-fold compared to non-inoculated plants (Pessina et al., 2014). Similarly, three out of four grapevine clade V *MLOs* (*VvMLO3*, *VvMLO4*, and *VvMLO17*) were induced during infection by *Erysiphe necator* (Feechan et al., 2008). Interestingly, in both apple and grapevine, a clade VII MLO was also found to respond to PM, but the significance of this finding is unclear. In our case, both clade V *CsMLOs* (*CsMLO1* and *CsMLO4*) appeared responsive to PM infection, showing a 2-fold upregulation (FDR < 0.01). This was further supported by the presence of *cis*-acting elements involved in gene overexpression by biotic and abiotic stresses in all of the clade V *CsMLOs* characterized in this study. These combined results suggest that cannabis is in a similar situation to that observed in *A. thaliana*, where all clade V genes could be involved in susceptibility.

As of now, *VrMLO12* in mungbean (*Vigna radiata*), which clusters with clade II genes, is the sole report of an *MLO* gene outside of clade V being clearly involved in PM susceptibility in dicots (Yundaeng et al., 2020). In rice, a clade II *MLO* (*OsMLO3*) was also found to have an expression pattern similar to clade V *MLOs* from *Arabidopsis* (Nguyen et al., 2016) and could partially restore PM susceptibility in barley mutants, suggesting an involvement in plant defense (Elliott et al., 2002). According to our results, clade II *CsMLOs* should not be considered candidates, as none showed an induction following inoculation with *G. ambrosiae* emend. (including *G. spadiceus*). However, other *CsMLOs* outside of clade V were found to be differentially expressed, a situation not different from previous findings in apple (Pessina et al., 2014), grapevine (Feechan et al., 2008), and tomato (Zheng et al., 2016). Interestingly, while only one out of four clade V *MLO* showed pathogen-dependent upregulation in tomato (*SlMLO1*), there is some overlap between non-clade V genes that respond to PM in tomato and cannabis. The pathogen-triggered response is, however, not always in the same direction. In tomato, three clade I *MLOs* (*SlMLO10*, *SlMLO13* and *SlMLO14*) were induced following an infection challenge with *Oidium neolycopersici*. While the cannabis orthologs of the last two (*CsMLO9* and *CsMLO2*, respectively) were also differentially expressed after inoculation with PM, their expression levels rather decreased over time (**Supplementary Figure S2**). Similarly, the expression of the tomato clade III *SlMLO4* significantly increased during infection, while that of its cannabis ortholog *CsMLO7* decreased (**Supplementary Figure S2**). The sole tomato clade VI gene, *SlMLO16*, was also found to be induced (Zheng et al., 2016). While its direct cannabis ortholog *CsMLO3* was not found to be up-regulated, it was the case for the other cannabis clade VI gene, *CsMLO5*, for which no ortholog exists in tomato or *Arabidopsis*. This particular gene was the only *CsMLOs* to be induced following a challenge with *G. ambrosiae* emend. (including *G. spadiceus*) outside of clade V.

Following our attempt to further characterize the two clade V CsMLOs and determine their subcellular localization *in planta* using confocal laser scanning microscopy, we observed that CsMLO4 was located in the plasma membrane while CsMLO1 was located in endomembrane-associated compartments, including the plasma membrane, the endoplasmic reticulum and the Golgi apparatus (**Figure 5**). In plants, the presence of a reticulate and network-looking pattern and bright spots, as observed for CsMLO1, are typically associated with the endoplasmic reticulum and Golgi stacks, respectively (Bassham et al., 2008). A time-series performed using confocal microscopy demonstrated that the CsMLO1-EGFP fusion protein was extremely dynamic compared to the CsMLO4-EGFP fusion protein (results not shown), supporting its implication in intracellular trafficking through the endomembrane system. Many studies have demonstrated that MLOs are localized in the plasma membrane (Devoto et al., 1999; Kim and Hwang, 2012; Nie et al., 2015), supporting our observations with regards to CsMLO4. Other studies have indicated that MLOs are associated with the plasma membrane and/or other endomembrane compartments, such as the endoplasmic reticulum and the Golgi apparatus (Chen et al., 2009; Jones and Kessler, 2017; Qin et al., 2019) and thus supporting our observations with regards to CsMLO1. To further support our findings and to determine precise subcellular localization of CsMLOs, subcellular fractionation studies as well as fluorescence colocalization with specific organelles markers should be performed.

### 4.4 Genetic variation within clade V *CsMLOs*: a quest toward durable resistance

Several natural or induced loss-of-function mutations have been identified in *MLO* genes that reduce susceptibility to PM (Büschges et al., 1997; Consonni et al., 2006; Pavan et al., 2011; Bai et al., 2008; Wang et al., 2014). Barley *HvMlo1* is probably the most studied *MLO* gene, and most mutations leading to loss-of-function (and thus, resistance to PM) in this gene appear to cluster in the second and third cytoplasmic loops (Reinstädler et al., 2010). The functional importance of those loops is still unclear, but evidence in other plants also points toward this particular region (Fujimura et al., 2016; Acevedo-Garcia et al., 2017). Outside of this region, the integrity of transmembrane domains, as well as certain invariant cysteine and proline residues, are critical for the function and accumulation of *MLO* proteins (Elliott et al., 2005; Reinstädler et al., 2010). Examination of polymorphisms among 32 cannabis cultivars (including one hemp variety) identified only a single SNP in the second cytoplasmic loop of *CsMLO1*. No mutations were identified among those invariant cysteine and proline residues, and no mutations were found either in any of the strictly conserved residues or within transmembrane domains. This suggests that loss-of-function mutations in *CsMLOs* could be rare or nonexistent among commercial cultivars, complicating future breeding efforts. The potential lack of diversity among the cultivars included in our analysis might also have impeded our ability to find causative loss-of-function mutations, which could be a scenario likely generalizable to a significant part of the cannabis production industry. Nevertheless, it is possible that mutations identified outside of those previously identified regions could inactivate *CsMLOs*. Three mutations causing amino acid replacements were identified in the first cytoplasmic loop of *CsMLO1*, and five mutations (four in *CsMLO1* and one in *CsMLO4*) were identified in the first extracellular loop. The proximal part of the C-terminus of MLO proteins contains a binding site for the cytoplasmic calcium sensor, calmodulin (Kim et al., 2002a; Kim et al., 2002b). In barley MLO, binding of calmodulin to this domain appears to be required for full susceptibility. Unfortunately, even though a high number of polymorphisms were identified in the cytoplasmic C-terminus of both *CsMLO1* (six mutations) and *CsMLO4* (10 mutations), those mutations are not found within the calmodulin-binding domain but rather at the distal end of the C-terminus. This might not be surprising, as this region is intrinsically disordered (Kusch et al., 2016). Intrinsically disordered regions, i.e. regions lacking stable secondary structures, usually exhibit greater amounts of non-synonymous mutations and other types of polymorphisms because of the lack of structural constraints (Nilsson et al., 2011; Kusch et al., 2016). Another study conducted on clade V *MLOs* had also revealed that both the first extracellular loop and the C-terminus were under strong positive selection (Iovieno et al., 2015), which seems in agreement with our observations.

The fact that potential loss-of-function mutations in clade V *MLOs* were not identified among the 32 investigated genomes suggests that complete resistance to PM might be hard to find among existing commercial cultivars. Nevertheless, such mutations might be present at a very low frequency, especially in “wild” populations or in landraces that have infrequently been used in breeding programs. For example, natural *mlo* alleles exist in barley but have only been found in landraces from Ethiopia and Yemen (Reinstädler et al., 2010). Similarly, the natural loss-of-function mutations in pea and tomato originated from wild accessions (Humphry et al., 2011; Bai et al., 2008). This should reinforce the importance of preserving cannabis wild populations and encourages efforts to establish germplasm repositories. However, *MLO*-based resistance to PM being a recessive trait, and assuming that both *CsMLO1* and *CsMLO4* are involved, this would mean that loss-of-function mutations would need to be bred as homozygous recessive for both genes into elite plants (not considering that multiple copies of a given *CsMLO* might exist, as suggested here for *CsMLO1*). In the absence of natural mutants, or to accelerate the implementation of *mlo*-based resistance in breeding programs, induced mutagenesis and genome editing might be interesting alternatives. While such approaches have not been optimized for cannabis, loss-of-function mutations have been obtained through those in other crops, such as barley (Reinstädler et al., 2010), wheat (Acevedo-Garcia et al., 2017; Wang et al., 2014), or tomato (Nekrasov et al., 2017).

There might also be other routes to combating PM than MLO. In crops where gene-for-gene interactions exist with PM (e.g., in cereals), a series of functional alleles confer complete resistance against distinct sets of PM isolates (Bourras et al., 2019). Since a similar situation has been observed between hop (*Humulus lupulus*, the closest relative of cannabis) and *Podosphaera macularis* (Henning et al., 2011), it is likely that such gene-for-gene interactions exist between cannabis and PM. In grapevine, resistance is usually considered polygenic, and there appear to be a diverse range of responses to invasion by *Erysiphe necator*, from penetration resistance to the induction of plant cell death (Feechan et al., 2011). Because cannabis has been excluded from scientific research for decades, very little is known about the biology of PM infection (i.e. genetic diversity, histology, host range), and no data on resistance levels among cultivars has been published as of now. Still, a recent study observed higher copy numbers of a thaumatin-like protein in several cannabis cultivars reported to be resistant to PM, and similar correlations were identified with copy number variations of endochitinases and *CsMLOs* (McKernan et al., 2020). Thaumatin-like proteins and endochitinases might thus represent additional targets potentially involved in quantitative resistance to PM.

## 5 Conclusion

Genome-wide characterization of *CsMLOs* performed in this study indicated that genes from clade V, which are immediately triggered upon infection by PM, are likely involved as early actors in the fungal infection process by the plant. Polymorphism data generated for clade V *CsMLOs* in multiple commercial cultivars suggested that loss-of-function mutations, required to achieve a resistance phenotype, are rare events and potentially challenging to assemble, especially when considering their recessive nature and the genetic redundancy of multiple clade V *CsMLOs*. Preserving a diversified collection of feral and, ideally, landrace cannabis genetic backgrounds while establishing coherent germplasm repositories could represent efficient strategies to compose with this complex biological reality. This will allow the creation of novel and valuable knowledge on the fundamental biology supporting the interaction between PM and cannabis, ultimately leading to more sustainable horticultural and agro-industrial practices.

## Author contributions

NP, FOH and DLJ contributed to the conception and design of the study. NP performed the experiments. NP, FOH and DLJ analyzed the data and drafted the manuscript. All authors revised the manuscript, read, and approved the submitted version. DLJ contributed reagents, equipment, and funds.

## Acknowledgments

We greatly thank François Sormany and Alex J. Cull for their help generating the RNA-seq and genomic datasets, respectively, Hugo Germain and his team for sharing their expertise with confocal microscopy, and Organigram for their continued support and for providing biomaterials.

## Data availability statement

All sequences generated in this study have been deposited in DDBJ/EMBL/GenBank under the BioProject accession: PRJNA738505, SRA accessions: SRR14839036-50; and the BioProject accession: PRJNA738519, SRA accessions: SRR14857079-110

## Supplementary materials

**Supplementary Table S1:** Members of the *CsMLO* gene family as predicted and manually curated in the CBDRx genome.

**Supplementary Table S2:** Members of the *CsMLO* gene family as predicted and manually curated in the Jamaican Lion (female) genome.

**Supplementary Table S3:** Members of the *CsMLO* gene family as predicted and manually curated in the Jamaican Lion (male) genome.

**Supplementary Table S4:** Members of the *CsMLO* gene family as predicted and manually curated in the Purple Kush genome.

**Supplementary Table S5:** Members of the *CsMLO* gene family as predicted and manually curated in the Finola genome.

**Supplementary Table S6:** Plant *cis*-acting regulatory elements identified in *CsMLOs* across five distinct cultivars.

**Supplementary Table S7:** Non-synonymous SNPs identified in *CsMLO1* and *CsMLO4* of 32 cannabis cultivars with their respective amino acid change in the resulting protein.

**Supplementary Figure S1:**
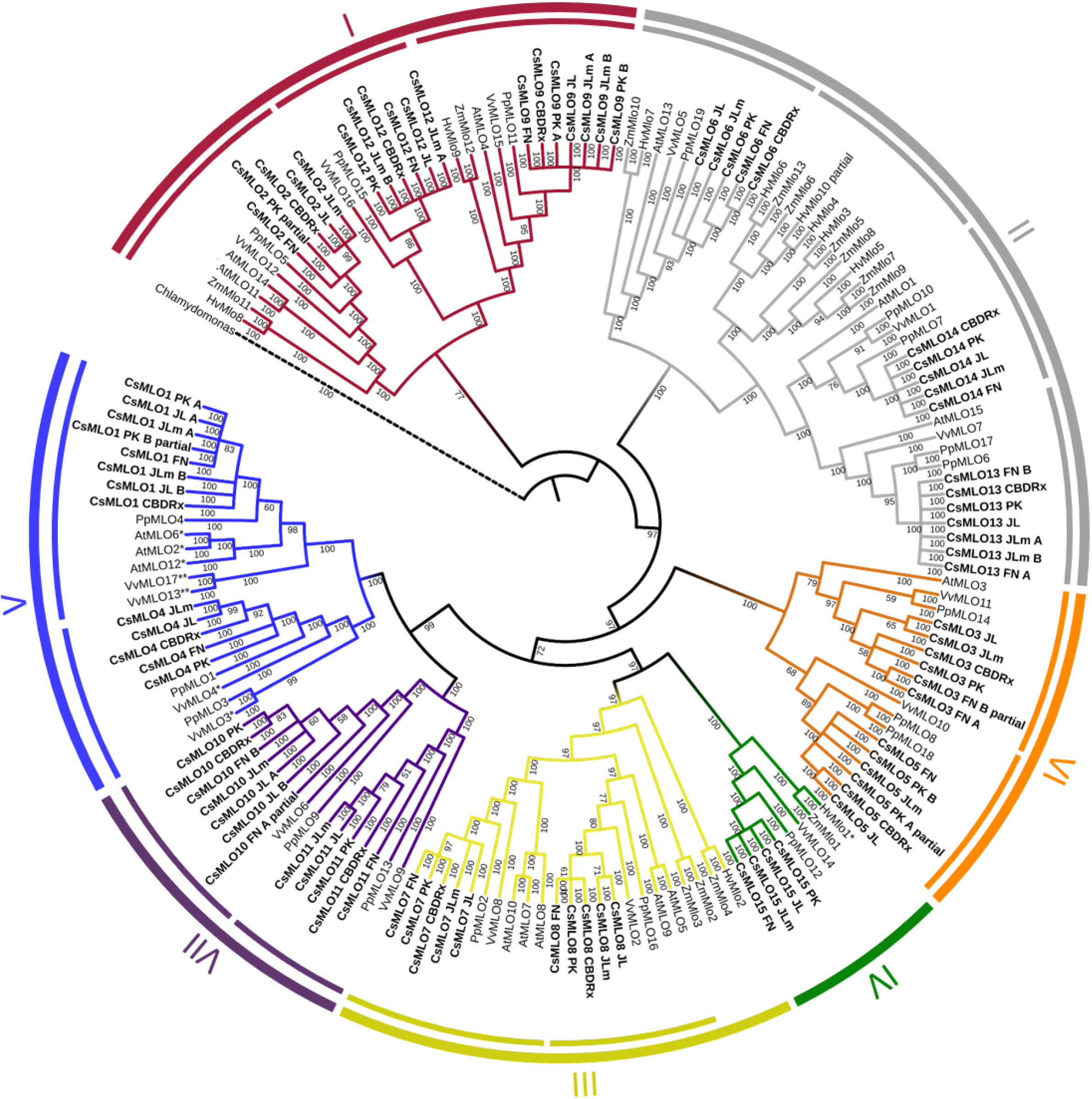
Phylogenetic relationships of CsMLOs based on Bayesian inference analysis. Phylogenetic tree of manually curated CsMLO proteins (bold) with MLO proteins from selected species (*Arabidopsis thaliana*, *Prunus persica*, *Vitis vinifera*, *Hordeum vulgare* and *Zea mays*). *Chlamydomonas reinhardtii* was used as an outgroup. Phylogenetic relationships were estimated using the MrBayes tool implemented on NGPhylogeny.fr, using default parameters. The seven defined clades are indicated, as well as potential subclades identified in this study (inner circles). Number on a node indicates the posterior probabilities of major clades and subclades. MLOs with one asterisk (*) have been experimentally demonstrated to be required for PM susceptibility (Büschges et al., 1997; Feechan et al., 2008; Wan et al., 2020), while MLOs with two asterisks (**) have been identified as main probable candidates for PM susceptibility (Pessina et al., 2016).

**Supplementary Figure S2.**
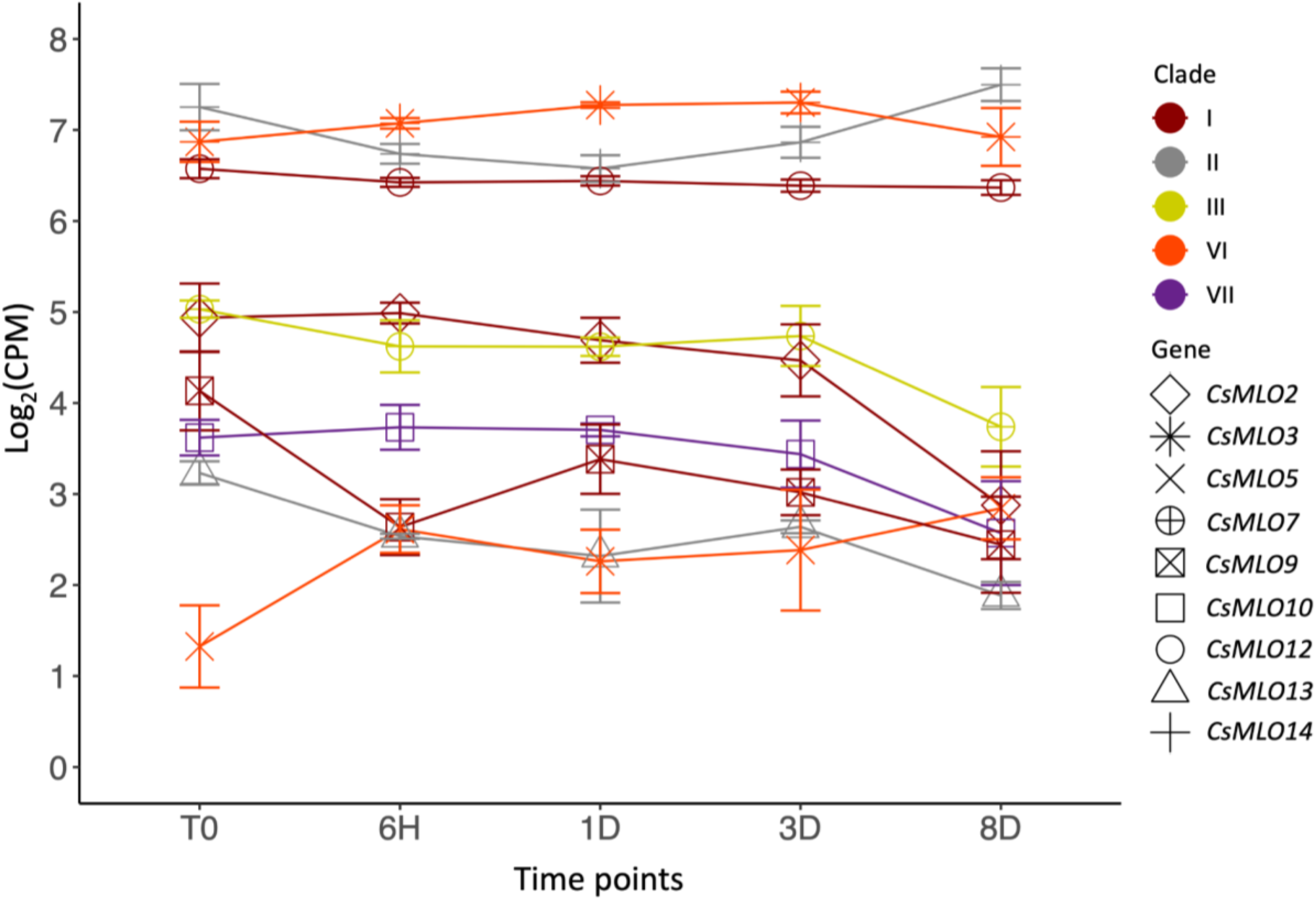
Transcriptomic response of *CsMLO* genes from all clades, except from clade V, following laboratory-controlled inoculations with powdery mildew. Specific *CsMLO* genes are represented by different symbols and the clade to which each gene belongs is displayed by different colors. Gene expression is displayed on the y-axis as the average logarithmic value of the Counts Per Million (logCPM) at each time point (displayed on the x-axis, n = 3 per time point). Time points: no infection / control (T0), 6 hours post-inoculation (6H), 24 hours post-inoculation (1D), 3 days post-inoculation (3D), and 8 days post-inoculation (8D).

**Supplementary File S1:** Nucleotide alignment of the genomic sequences from all 86 *CsMLOs*. Exons and introns are separated by multiple gaps in the alignment, as is the case for the insertion in the first exon of *CsMLO5-JLm*.

**Supplementary File S2:** Nucleotide alignment of the coding sequences from all 86 *CsMLOs*.

**Supplementary File S3:** Amino acid alignment of the protein sequences from all 86 *CsMLOs*.

